# Decoupling axonal regrowth and branching through Imp-dependent RNA regulation during neuronal remodeling

**DOI:** 10.64898/2026.07.27.740905

**Authors:** Margot Noguères, Zeinab Rekad, Florence Besse, Caroline Medioni

## Abstract

Structural remodeling of neuronal projections in response to developmental cues, injury, or disease is essential for adaptive circuit rewiring. This dynamic process, characterized by pruning and regrowth phases, requires the coordinated execution of neurite regrowth and branching to establish functional neuronal circuits. Yet, how these processes are regulated in space and time at the post-transcriptional level remains poorly understood. Here, we identify the conserved RNA-binding protein Imp (IGF2BP) as a central regulator of developmental axonal remodeling in *Drosophila* CCAP/Bursicon neurons. We show that Imp acts within a restricted time window during late metamorphosis to control both late regrowth and branching of adult CCAP/Bursicon axons. Combining functional approaches, high-resolution imaging and single-molecule mRNA detection, we further show that Imp controls these temporally distinct programs through genetically separable regulatory mechanisms. While axonal elongation is mediated by Imp-dependent stabilization of *profilin* mRNA, axonal branching is mediated by an independent mechanism that may involve local regulation in axons. Together, our findings demonstrate that axonal regrowth and branching, two morphogenetic events essential for neuronal circuit maturation *in vivo*, are controlled independently, yet coordinated through a common and conserved post-transcriptional framework.

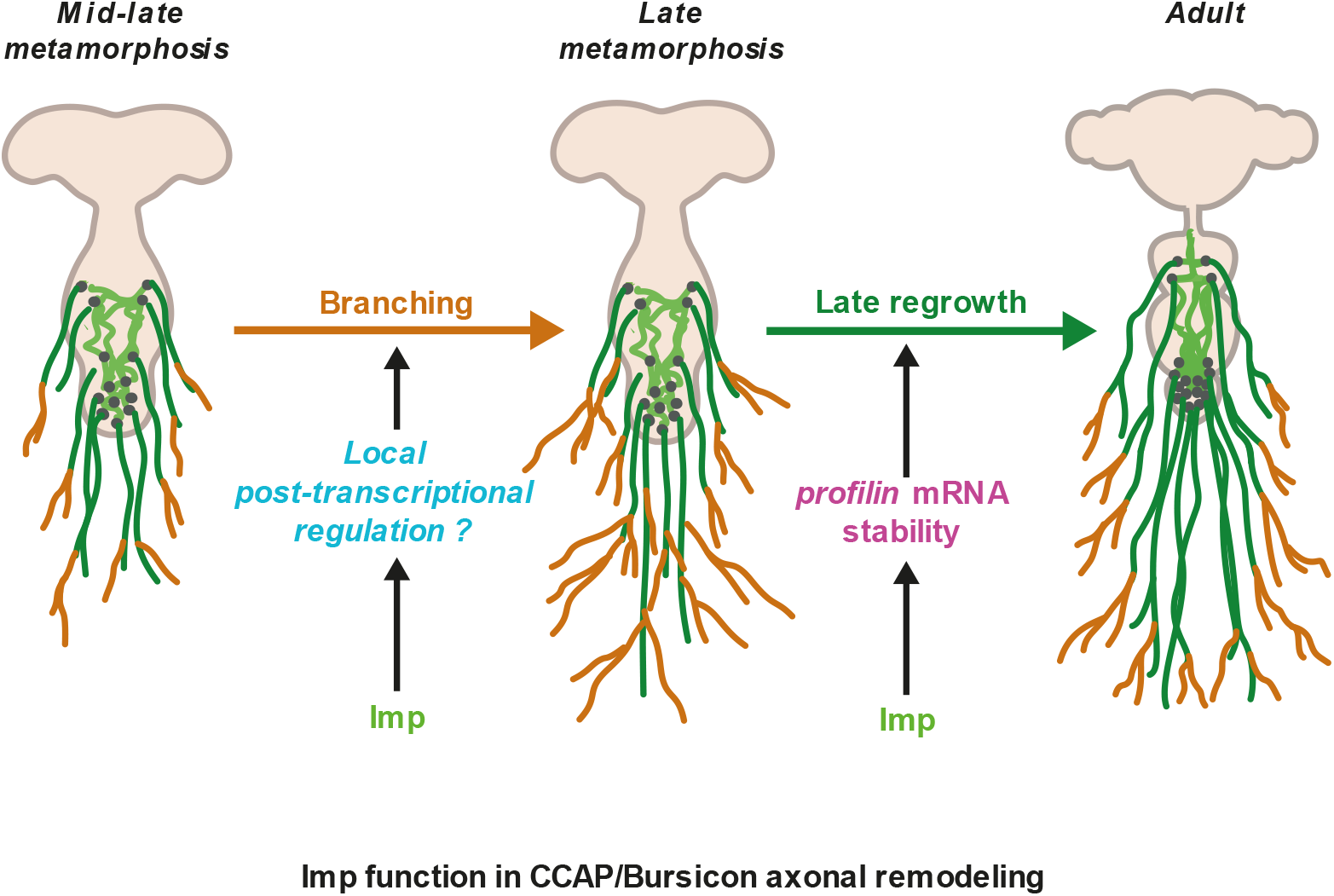

## Introduction

The establishment of functional neural circuits requires not only the generation of appropriate neuronal identities but also the precise remodeling of neuronal processes throughout development. Diverse neuronal populations undergo extensive structural reorganization involving axonal pruning followed by regrowth and branching to establish mature connectivity patterns. Such remodeling events are particularly prominent during developmental transitions, including insect metamorphosis or vertebrate postnatal development, where larval or immature circuits are transformed into adult functional networks (Luo and O’Leary 2005; Yaniv and Schuldiner 2016). Although substantial progress has been made in identifying extracellular signals and transcriptional programs controlling neuronal remodeling (Riccomagno and Kolodkin 2015; Alyagor et al. 2018; Meltzer and Schuldiner 2022), the post-transcriptional mechanisms coordinating these complex morphogenetic processes remain incompletely understood.

RNA-binding proteins (RBPs) play essential roles in neuronal development by regulating mRNA stability, localization and translation to meet the differential requirements of soma and distant axonal and dendritic compartments (Jung et al. 2012; Holt et al. 2019). Through the assembly of ribonucleoprotein (RNP) granules and the transport of target transcripts, RBPs control diverse aspects of neuronal morphogenesis, including axon guidance, branching, synaptogenesis, and regeneration (Darnell 2013; Turner-Bridger et al. 2020). However, how individual RBPs coordinate distinct phases of developmental neuronal remodeling remains poorly understood.

Members of the conserved Insulin-like Growth Factor 2 mRNA-Binding Protein (IGF2BP) family of RBPs have emerged as major regulators of RNA localization and local translation in the nervous system. In vertebrates, IGF2BP1 (also known as ZBP1) controls the transport and translation of *β-actin* mRNA in growth cones and contributes to axon growth and regeneration *in vitro* and *in vivo* (Zhang et al. 2001; Leung et al. 2006; Yao et al. 2006; Donnelly et al. 2011). In *Drosophila,* Imp, the sole orthologue of the vertebrate IGF2BP family, assembles into RNP particles that are selectively transported to the axons of central nervous system neurons during development, suggesting a conserved function in neuronal morphogenesis. Indeed, in Mushroom Body γ neurons, Imp was shown to promote the regrowth of adult axonal projections following pruning during metamorphosis, a process that requires the regulation of *profilin* mRNA (Medioni et al. 2014). Whether Imp employs similar mechanisms in other remodeling neuronal populations, and how its functions are spatially and temporally organized to coordinate distinct morphogenetic events during neuronal remodeling, remain largely unknown. The *Drosophila* CCAP/Bursicon neurons provide a tractable system to investigate how post-transcriptional mechanisms coordinate distinct phases of neuronal remodeling *in vivo*. During pupal development, CCAP/Bursicon neurons eliminate larval projections and subsequently regrow complex adult axonal arbors to establish a projection network required for wing expansion after eclosion (Luan et al. 2006; Peabody et al. 2008). Defects in CCAP/Bursicon neuronal remodeling have been associated with impaired wing expansion, underscoring the functional importance of this remodeling process (Zhao et al. 2008). Although several transcriptional and hormonal pathways regulating CCAP/Bursicon neuron development have been identified (Zhao et al. 2008; Gu et al. 2014; Yaniv and Schuldiner 2016; Chen et al. 2017; Gu et al. 2017), the contribution of post-transcriptional regulation to CCAP/Bursicon neuronal remodeling remains poorly characterized.

Here, we identify Imp as a key regulator of CCAP/Bursicon neuron axonal remodeling. We show that Imp is specifically required during a restricted time-window at late metamorphosis to promote the establishment of the adult axonal network. Imp regulates late axonal regrowth and branch formation through separable mechanisms. Indeed, we demonstrate that regulation of *profilin* mRNA stability underlies Imp-dependent axonal regrowth but not branching, revealing a molecular uncoupling between these two processes. Finally, our analysis suggest that the axonal localization of Imp may contribute specifically to branch formation. Together, our results support a model in which temporally restricted post-transcriptional regulation coordinates distinct aspects of neuronal remodeling.

## Results

### Imp controls the axonal morphology and function of adult CCAP/Bursicon neurons

To investigate if Imp is required for CCAP/Bursicon axonal remodeling, we inactivated *imp* using the CCAP-Gal4-driven expression of a deGradFP construct (Caussinus and Affolter 2016) combined with *gfp*-RNAi in GFP-Imp knock-in flies (hereafter referred to as *imp* Degrad condition). CCAP/Bursicon neuron projections extend from the Ventral Nerve Cord (VNC) to form a stereotyped and highly elaborate arborized network throughout the abdominal cavity of adult *Drosophila* (Figure 1A), which we visualized using either CCAP-Gal4-driven expression of mCD8-RFP or anti-Bursicon immunolabeling. Both methods produced identical axonal labeling patterns (Figure S1A,B).

**Figure 1:**
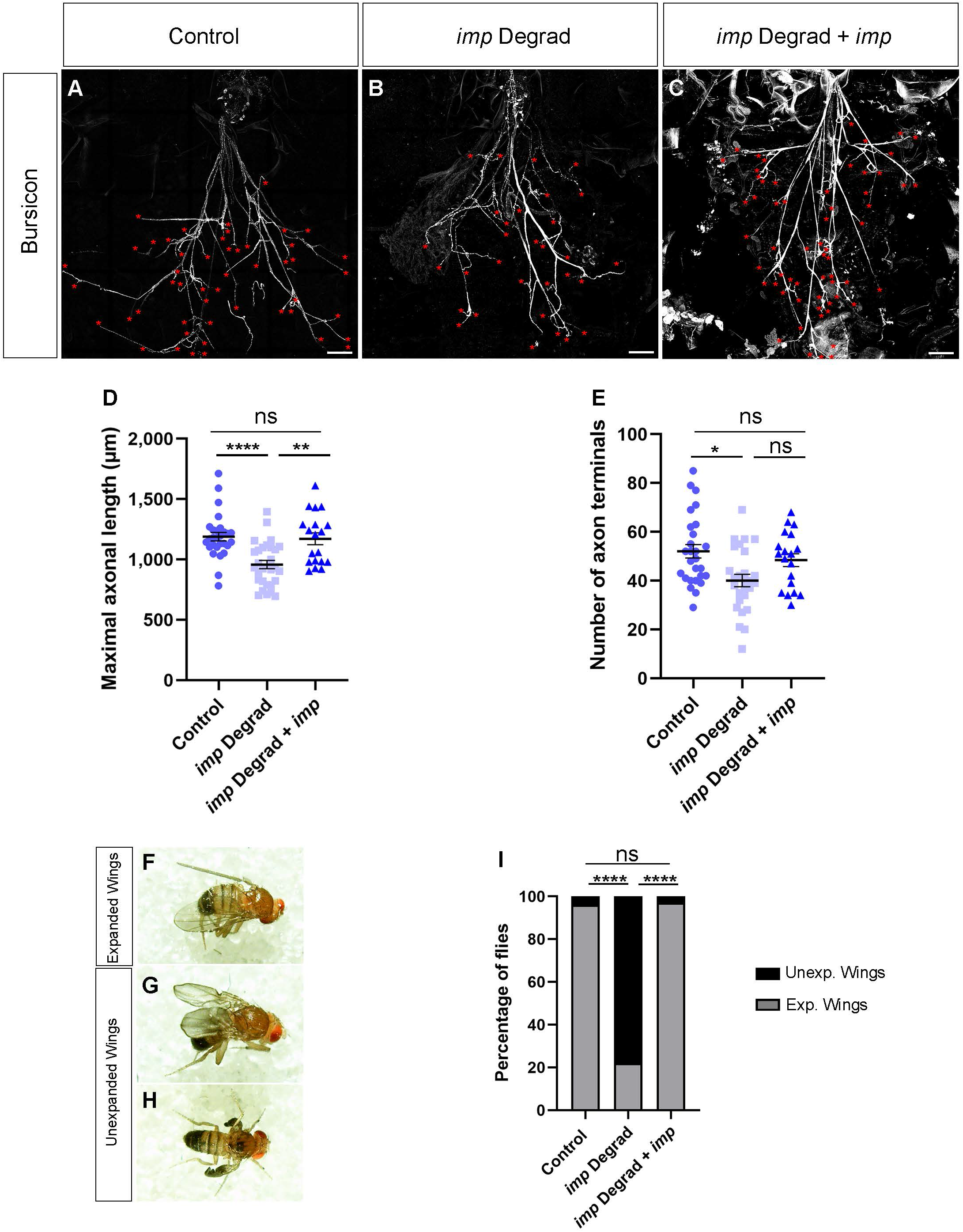
*imp* inactivation induces elongation and branching defects of CCAP/Bursicon axons and wing expansion defects. **A-C** Tile confocal images of control (**A**), *imp* Degrad (**B**) and *imp* Degrad+*imp* (**C**) obtained from whole-mount fillets dissected 24 h post-eclosion and stained using anti-Bursicon antibody. Scale bar: 100 µm. Red asterisks indicate the axon terminals. Animals were raised at 29°C. Complete genotypes: control: GFP-Imp/Y; CCAP-Gal4,UAS-mCD8-RFP/+; *imp* Degrad: GFP-Imp/Y; CCAP-Gal4,UAS-mCD8-RFP/+; UAS-*degrad-fp*,UAS-*gfp*-RNAi/+; *imp* Degrad+*imp*: GFP-Imp/Y; CCAP-Gal4/+; UAS-*degrad-fp*,UAS-*gfp*-RNAi/ UASp-*flag*-*imp*. **D** Dot plot showing the maximal axonal length of control, *imp* Degrad and *imp* Degrad+*imp* CCAP/Bursicon neurons at 24 h post-eclosion. Number of fillets analyzed: control=28; *imp* Degrad=29; *imp* Degrad+*imp*=19. **E** Dot plot showing the number of CCAP/Bursicon axon terminals in control, *imp* Degrad and *imp* Degrad+*imp* adult flies at 24 h post-eclosion. Number of fillets analyzed: control=27; *imp* Degrad=27; *imp* Degrad+*imp*=19. **D-E** 2 to 5 replicates were performed. Quantifications were performed on the same individuals when possible. Bars and error bars correspond respectively to mean and SEM. ****P<0.0001; **P<0.01; *P<0.05 (Kruskall–Wallis with Dunn’s post-tests). **F-H** Example of expanded (**F**) and unexpanded (**G-H**) wing phenotypes. **I** Bar graph showing the frequency of flies with expanded wings (Exp. Wings, gray) and unexpanded wings (Unexp. Wings, black) in control, *imp* Degrad and *imp* Degrad+*imp* conditions. 4 replicates were performed and the proportion of each wing phenotype was calculated from the cumulative number of flies. Number of individuals scored per condition: control=1,699; *imp* Degrad=2,006; *imp* Degrad+*imp*=1,045. ****P<0.0001 (Chi-squared (χ²) test).

Quantitative analysis revealed a significant reduction in both the length of axons (Figure 1B,D) and the number of axon terminals (Figure 1B,E) following *imp* inactivation, arguing that Imp normally promotes axonal elongation and branching of CCAP/Bursicon neurons.

To rule out the possibility that *imp* inactivation induces differentiation defects, we analyzed two cellular parameters characteristic of mature CCAP/Bursicon neurons in adult flies: CCAP/Bursicon cell body number and size. No significant changes were observed in cell body number or size following *imp* inactivation (Figure S2A-D), indicating that Imp acts specifically on axonal morphology independently of differentiation.

To confirm the specificity of Imp requirement in axon elongation and branching, we re-introduced *imp* expression in CCAP/Bursicon neurons in the *imp* Degrad loss-of-function background. This fully restored maximal axonal length (Figure 1C,D) but rescued partially the number of axon terminals (Figure 1C,E), suggesting that branching is more sensitive to *imp* function than regrowth.

Given the known function of adult CCAP/Bursicon neurons in promoting wing expansion after eclosion (Luan et al. 2006; Peabody et al. 2008; Zhao et al. 2008), we next assessed whether the severe axonal arborization defects induced by *imp* inactivation may result in wing expansion phenotype. *imp* inactivation caused wing expansion defects in more than 70% of adult *Drosophila* (Figure 1F-I). These phenotypes were fully reverted by re-expression of *imp in* CCAP/Bursicon neurons (Figure 1I).

Together, these results demonstrate that Imp cell-autonomously controls both axonal elongation and branching of CCAP/Bursicon neurons, thereby ensuring their proper physiological function.

### Imp acts during late metamorphosis to promote the functional maturation of adult CCAP/Bursicon neurons

CCAP/Bursicon neurons are born during embryonic development, then undergo extensive remodeling during metamorphosis (Luan et al. 2006; Zhao et al. 2008). Thus, we sought to define the critical period during which *imp* function is required in CCAP/Bursicon neurons. To this end, we conditionally inactivated *imp* at defined stages of development and used the wing expansion phenotype as a simple and robust readout of CCAP/Bursicon axonal complexity.

To achieve temporal control of *imp* inactivation, we took advantage of the well-characterized temperature sensitivity of the Gal4 protein (Nagarkar-Jaiswal et al. 2015), whose transcriptional activity is highly attenuated at 18 °C compared to 25 °C. While at 25°C, deGradFP and *gfp-*RNAi are highly expressed and *imp* thus significantly downregulated, deGradFP and *gfp-*RNAi are poorly expressed at 18°C and *imp* expression preserved.

Shifting *imp* Degrad flies from 18°C to 25 °C for the full duration of metamorphosis recapitulated the penetrance of the wing expansion phenotype observed upon constitutive *imp* inactivation (Figure 2A,B, top). Conversely, shifting *imp* Degrad flies from 25°C to 18 °C for the full duration of metamorphosis had only minimal effect on wing expansion (Figure 2A, bottom). These results thus indicate that Imp is required during metamorphosis to promote adult CCAP/Bursicon neuron function.

**Figure 2:**
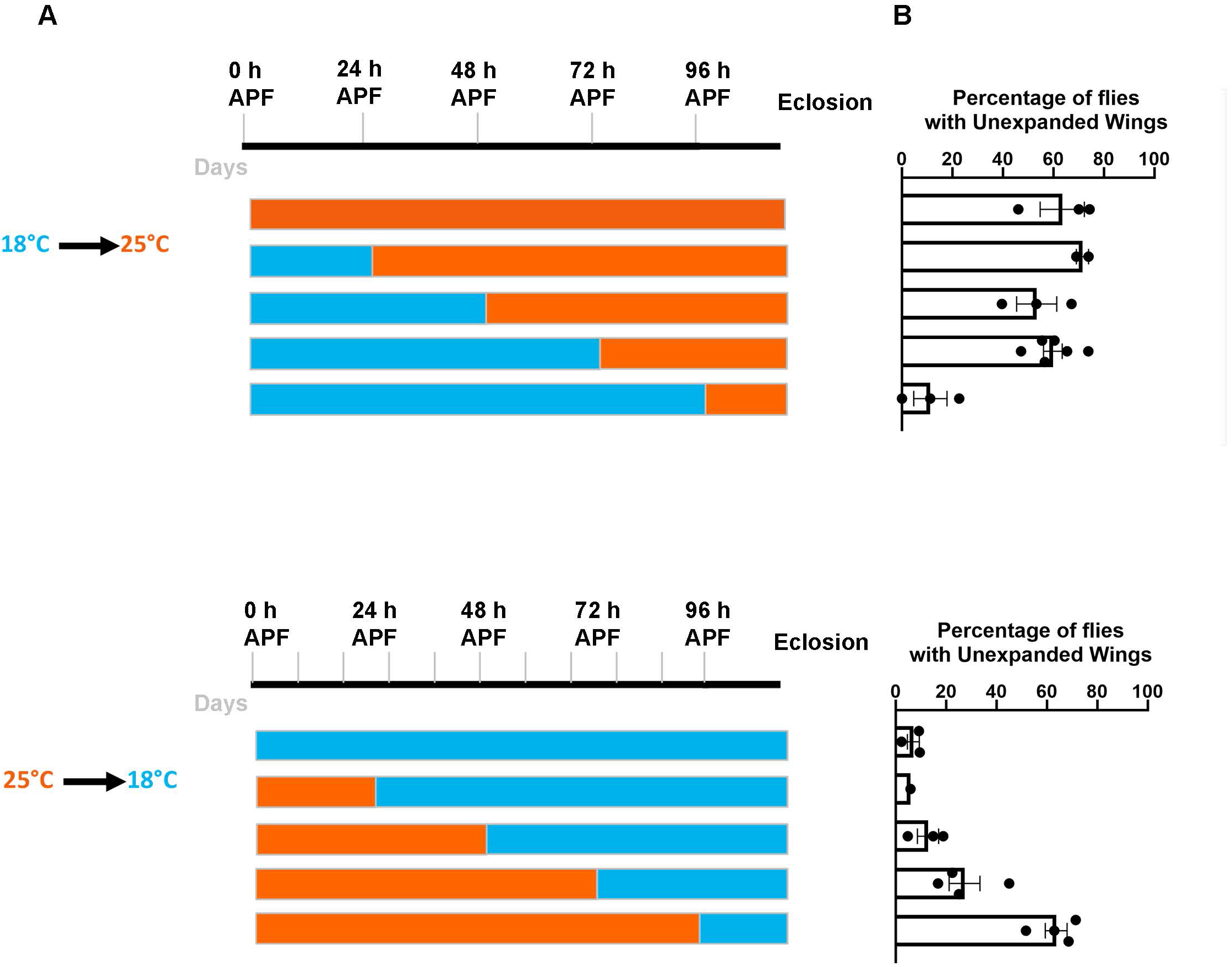
*imp* function is required at late metamorphosis to control wing expansion. **A** Diagrams illustrating the temperature shifts performed to inactivate *imp* specifically in CCAP/Bursicon neurons. The top black line indicates developmental stages from 0 hour After Puparium Formation (h APF) to eclosion, with correspondence in days indicated in gray for animals reared at 25°C (top) and 18°C (bottom). Bars below indicate the time-points at which the animals were shifted from 18°C (blue, *imp* Degrad OFF) to 25°C (orange, *imp* Degrad ON) (top), or from 25°C to 18°C (bottom). Complete genotypes: control: GFP-Imp/Y; CCAP-Gal4,UAS-mCD8-RFP/Cyo; *imp Degrad*: GFP-Imp/Y; CCAP-Gal4,UAS-mCD8-RFP/+; UAS-*degrad-fp*,UAS-*gfp*-RNAi/+. **B** Bar graphs showing the proportion of flies with unexpanded wings in the different conditions described in **A**. Replicate experiments (from 1 to 6 replicates per condition) are represented as individual data points. Bars and error bars represent respectively the mean proportion of unexpanded wings in the different replicates and SEM. Number of individuals per condition for shifts from 18 to 25 °C (top): from 0 h APF to adult=899; 24 h APF to adult=714; 48 h APF to adult=815; 72 h APF to adult=1,542; 96 h APF to adult=581. Number of individuals per condition for shifts from 25 to 18 °C (bottom): from 0 h APF to adult=712; 24 h APF to adult=231; 48 h APF to adult=877; 72 h APF to adult=1,354; 96 h APF to adult=457.

To further refine the temporal requirement of *imp* function, we then restricted *imp* inactivation to distinct developmental windows by shifting *imp* Degrad flies from 18°C to 25°C from 24 hours After Puparium Formation (h APF) to eclosion, 48 h APF to eclosion, 72 h APF to eclosion or 96 h APF to eclosion (Figure 2A, top). Wing expansion defects were observed in all conditions but the last one, indicating that Imp activity is specifically required during the latest phase of metamorphosis (Figure 2A,B, top). Consistent with this, reverse shift experiments in which *imp* function is inactivated until 48 h APF had no impact on wing expansion, while wing expansion defects were observed upon shifting flies to 18°C at 72 h APF or 96 h APF (Figure 2A,B, bottom).

Given the expected delay between temperature shifts and the resulting changes in Imp levels, these results identified a specific window during late metamorphosis — between 72 and 96 h APF — as the critical period during which Imp activity is essential for the maturation of CCAP/Bursicon neurons.

### Imp-dependent axonal branching and regrowth are temporally uncoupled

While our conditional inactivation experiments identified when *imp* is required, they did not precisely address how *imp* regulates the temporal sequence of axon elongation and branching during metamorphosis.

To address this, we first quantified axonal length and number of axon terminals in control conditions at successive stages of metamorphosis. In control, larval CCAP/Bursicon axons extended through the abdominal cavity with a mean maximal axonal length of ∼1,100 µm (Figure 3A-C) and displayed claw-like structures previously described as neuroendocrine endings (Hodge et al. 2005; Vomel and Wegener 2007; Zhao et al. 2008). Axonal pruning had already started at 14 h APF and was complete at 30 h APF, when most individuals had no axonal projections detectable outside the VNC (Figure 3A-C). Axonal regrowth initiated around 48 h APF, with short axonal processes extending posteriorly (Figure 3A-C). Axons reached half of their adult length by 72 h APF (Figure 3A-C) and continued to elongate until adult stage, to reach a mean maximal axonal length of ∼1,200 µm 24 h post-eclosion (Figure 3A-C). Quantitative temporal analysis revealed that, whereas pruning proceeded as a single uniform phase, regrowth was bi-phasic, characterized by a rapid initial regrowth phase between 48 h and 72 h APF (∼600 µm/day) followed by a slower late regrowth phase from 72 h APF to adult stage (∼200µm/day).

**Figure 3:**
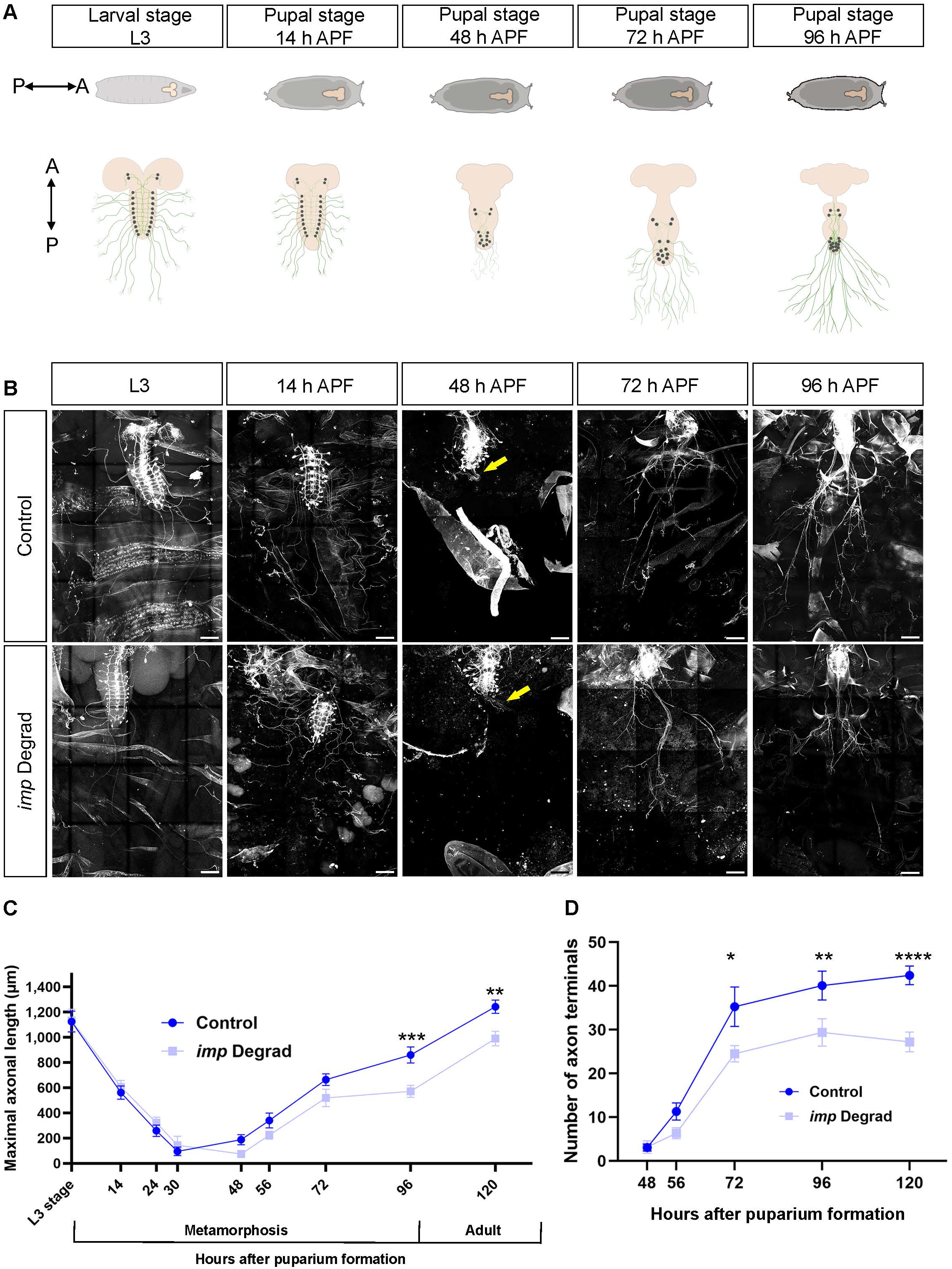
*imp* inactivation disrupts two temporally distinct phases of CCAP/Bursicon axonal remodeling: branching and late regrowth. **A** Schematic representation of the ventral nerve cord (top, in beige) and of CCAP/Bursicon neuron axonal remodeling (bottom, in green) at different stages, from wandering third-instar larval stage (L3) to 96 h APF. **B** Tile confocal images of control (upper row) and *imp* Degrad (lower row) obtained from whole-mount fillets at L3, 14 h, 48 h, 72 h and 96 h APF. CCAP/Bursicon neurons were labeled using CCAP-Gal4 driven mCD8-RFP expression. At 48 h APF, Fredllow arrows highlight the fine axonal projections. Scale bar: 100 µm. Animals were raised at 25°C. Complete genotypes: control: GFP-Imp/Y; CCAP-Gal4,UAS-mCD8-RFP/CyO; *imp* Degrad: GFP-Imp/Y; CCAP-Gal4,UAS-mCD8-RFP/+; UAS-*degrad-fp*,UAS-*gfp*-RNAi/+. **C** Time course analysis of maximal axonal lengths from L3 to adult stage in control (dark blue) and *imp* Degrad (light blue) conditions. Number of fillets analyzed for each time-point in control condition: L3=12, 14 h APF=15, 24 h APF=15, 30 h APF=20, 48 h APF=18, 56 h APF=14, 72 h APF=20, 96 h APF=29, Adult=25; *imp* Degrad condition: L3=14, 14 h APF=12, 24 h APF=11, 30 h APF=9, 48 h APF=10, 56 h APF=30, 72 h APF=12, 96 h APF=23, Adult=23. **D** Time course analysis of the number of axon terminals from 48 h APF to adult stage in control (dark blue) and *imp* Degrad (light blue) conditions. Number of fillets analyzed for each time-point in control condition: 48 h APF=21, 56 h APF=7, 72 h APF=20, 96 h APF=28; Adult=24; in *imp* Degrad condition: 48 h APF=10, 56 h APF=15, 72 h APF=9, 96 h APF=18, Adult=17. **C-D** Quantifications were performed on the same individuals when possible. Data points correspond to mean values and error bars to SEM. ****P<0.0001; ***P<0.001; **P<0.01; *P<0.05 (two-tailed Mann-Whitney U test on individual data points).

Axonal branching followed a distinct temporal profile. Ramification of axonal arbors initiated by 56 h APF and peaked at 72 h APF, with a number of axon terminals increasing from ∼10 to ∼35 during this temporal window (Figure 3A,B,D). Minimal addition of branches was observed from 72 h APF to adulthood, when a mean of ∼42 axon terminals was observed (Figure 3A,B,D). Thus, branching occurs mainly during the fast initial regrowth phase, whereas minimal additional branching occurs during the late regrowth phase.

Consistent with *imp* temporal requirement, CCAP-specific *imp* inactivation did not induce detectable alterations in pruning or initial regrowth until 72 h APF (Figure 3B-D). Axonal length defects were observed from 96 h APF, reflecting a specific impairment of the late regrowth phase characterized by a 4-fold reduction of regrowth rate (∼50 µm/day in *imp* Degrad flies versus ∼200 µm/day in controls; Figure 3B,C). In contrast, branching defects were already detectable at 72 h APF, with a ∼30% decrease in the number of axon terminals (∼25 in *imp* Degrad flies versus ∼35 in controls, Figure 3B,D).

Together, these results indicate that Imp controls two processes with distinct temporality: the branching of CCAP/Bursicon axons and their subsequent late phase of regrowth.

### Late axonal regrowth coincides with increased Imp levels

Having identified a specific temporal requirement for Imp function during late metamorphosis, we wondered if Imp levels or subcellular distribution were regulated at this stage.

Imp being a conserved component of neuronal RNP granules (Tiruchinapalli et al. 2003; Leung et al. 2006; Yao et al. 2006), we first characterized its subcellular distribution in CCAP/Bursicon cell bodies through high-resolution confocal imaging at 48, 72 and 96 h APF (Figure 4A-D). At all stages analyzed, Imp displayed a dual distribution, localizing both in a diffuse cytoplasmic pool and in granular structures. GFP-Imp granule density remained constant across metamorphosis in control animals, but was significantly reduced following *imp* inactivation (Figure 4E), confirming the specificity of the granular signal observed in CCAP/Bursicon cell bodies.

**Figure 4:**
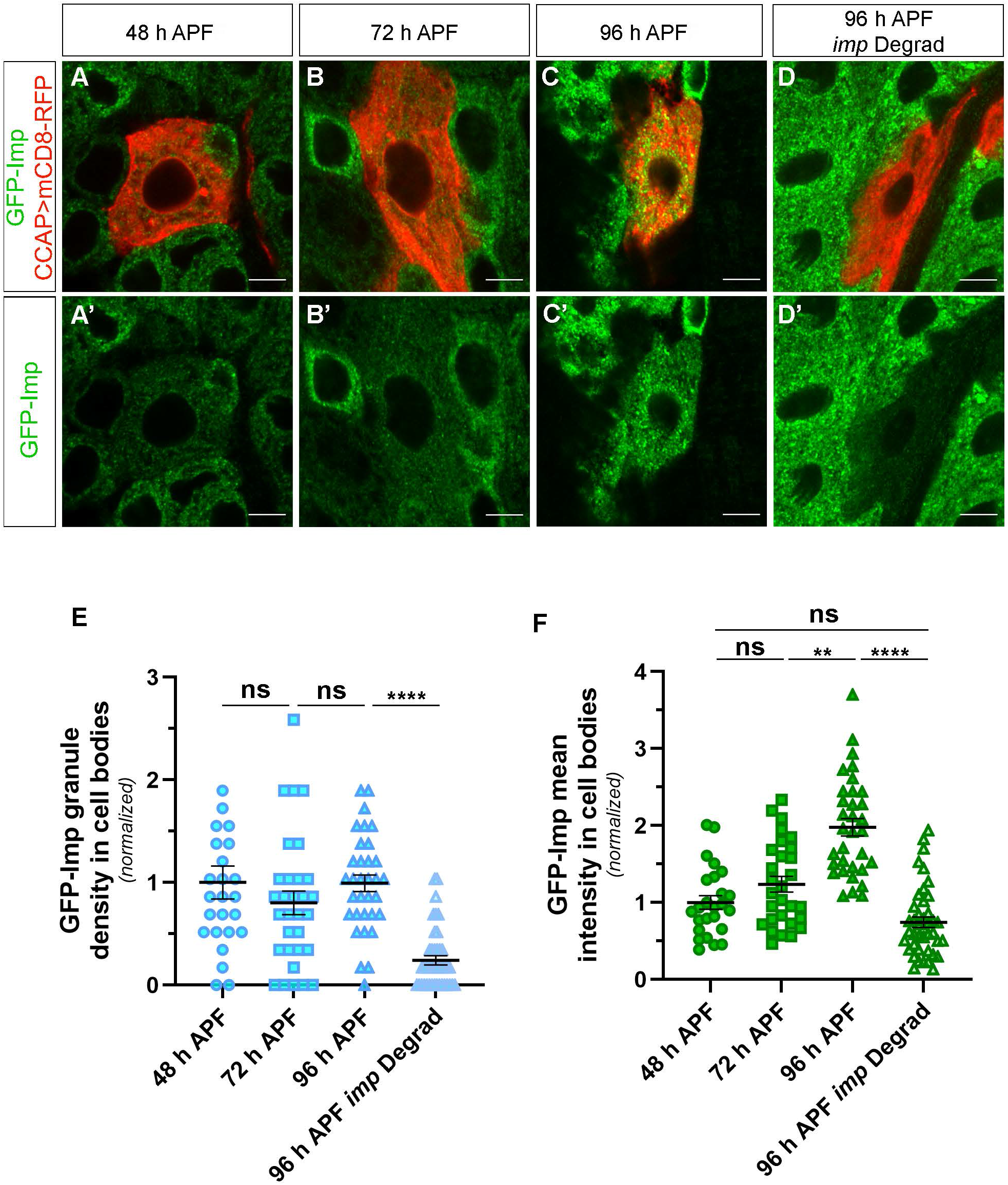
Late axonal regrowth is associated with increased Imp levels but unchanged granule density. **A-D** Confocal images of CCAP/Bursicon neuron cell bodies obtained from control (**A-C**) or *imp* Degrad (**D**) whole-mount fillets at 48 h APF (**A**), 72 h APF (**B**), 96 h APF (**C,D**). CCAP/Bursicon neurons were detected using CCAP-Gal4 driven mCD8-RFP expression (in red in **A-D**), and GFP-Imp signal is shown in green in **A-D** and **A’-D’**. Scale bar: 5 µm. Animals were raised at 25°C. Complete genotypes: control: GFP-Imp/Y; CCAP-Gal4,UAS-mCD8-RFP/CyO; *imp* Degrad: GFP-Imp/Y; CCAP-Gal4,UAS-mCD8-RFP/+; UAS-*degrad-fp*,UAS-*gfp*-RNAi/+. **E** Dot plot showing the GFP-Imp granules in CCAP cell bodies at 48 h APF, 72 h APF, 96 h APF, and 96 h APF in the *imp* Degrad condition. **F** Dot plot showing GFP-Imp mean intensity in CCAP/Bursicon cell bodies at 48 h APF, 72 h APF, 96 h APF, and 96 h APF in the *imp* Degrad condition. **E-F** Number of CCAP/Bursicon cell body analyzed: 48 h APF=25, 72 h APF=29, 96 h APF=33, and 96 h APF *imp* Degrad=42. 2 to 3 replicates were performed. Quantifications were performed on the same individuals. Individual data points correspond to individual cell bodies for each condition and were normalized to the 48 h APF condition. 1 to 6 CCAP/Bursicon cell bodies were analyzed per individual. Bars and error bars correspond respectively to mean and SEM. ****P<0.0001; **P<0.01; ns stands for not significant (Kruskall–Wallis with Dunn’s post-tests).

In parallel to our analysis of somatic Imp subcellular distribution, we monitored Imp protein levels in CCAP/Bursicon cell bodies by quantifying endogenous GFP-Imp fluorescence intensity (Figure 4A-D). This revealed that Imp expression levels remained stable between 48 h and 72 h APF, but increased approximately two-fold between 72 and 96 h APF (Figure 4F). As a proof of specificity, GFP-Imp levels were significantly decreased under *imp* Degrad condition. These results thus identify a stage-specific increase in Imp abundance that coincides with Imp function in late axonal regrowth.

### Imp-mediated axonal branching and late regrowth can be genetically uncoupled

Given the established ability of Imp to localize within axonal processes (Medioni et al. 2014; Vijayakumar et al. 2019), we wondered whether Imp axonal localization contributes to axonal morphogenesis. First, we asked whether Imp was present within CCAP/Bursicon axons, thereby defining a potential axonal pool that could contribute to local control of axonal morphogenesis. Consistent with this possibility, GFP-Imp was readily detected in granules within distal axonal regions during late metamorphosis (72 and 96 h APF) and remained detectable at the adult stage (Figure S3A-B,5A).

Then, we took advantage of the constitutive *imp* ΔPLD mutant, which was previously shown to reduce Imp axonal transport in Mushroom Body γ neurons (Vijayakumar et al. 2019). In this CRISPR mutant, a GFP-tagged form of Imp deleted of its C-terminal low-complexity prion-like domain (PLD) is expressed from the endogenous locus. As expected, *imp* ΔPLD mutant flies showed a significant reduction in GFP-Imp levels within CCAP/Bursicon axons compared to controls (Figure 5A-C). In addition, a decrease in GFP-Imp levels was also detected in *imp* ΔPLD CCAP/Bursicon cell bodies (Figure S3C-F). Importantly, this reduction was substantially less pronounced than that observed under *imp* Degrad condition (Figure S3D-F).

**Figure 5:**
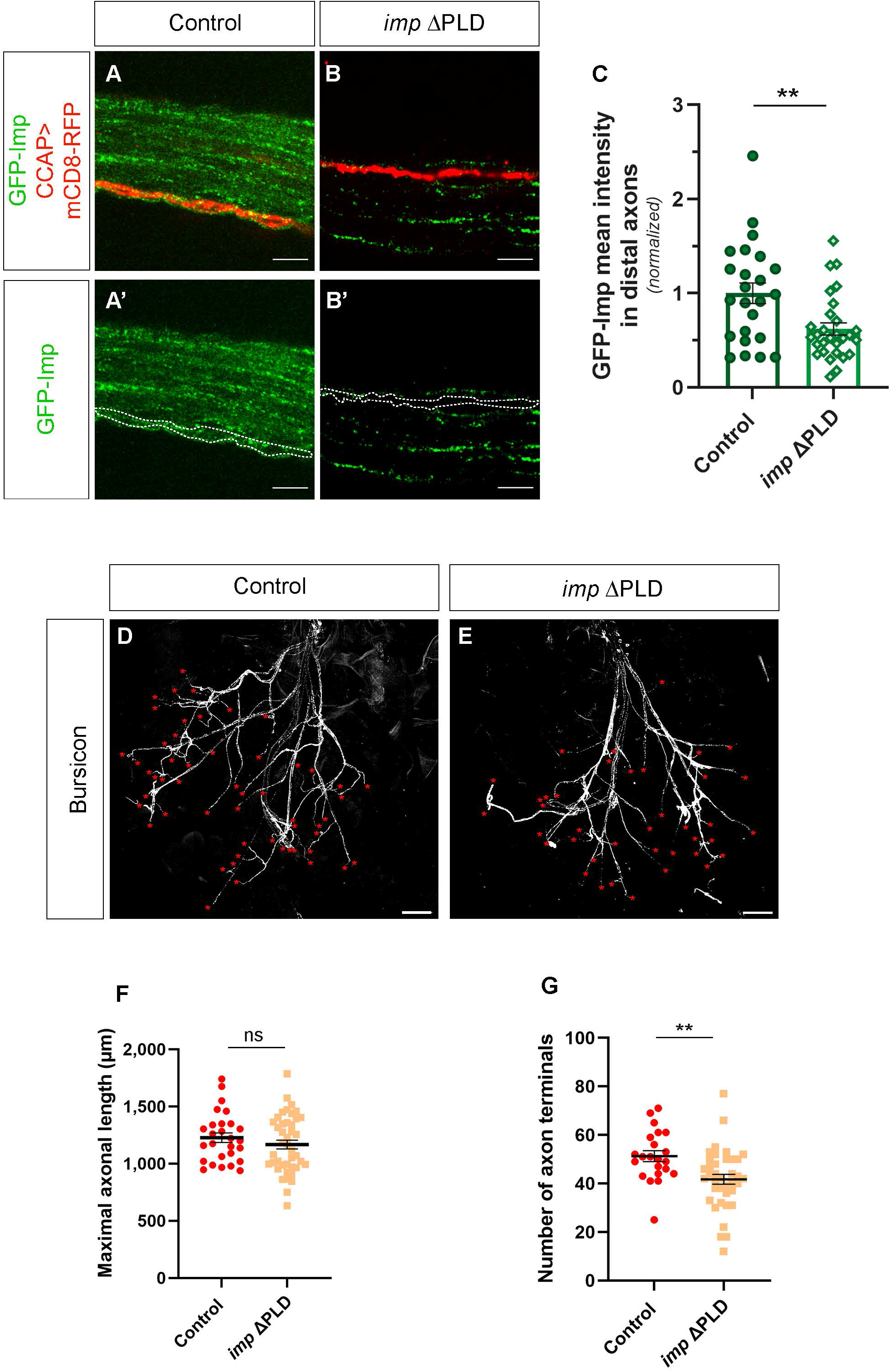
Axonal branching, but not late regrowth, is altered in *imp* ΔPLD mutants. **A-B** Confocal images of CCAP/Bursicon neuron distal axons obtained from control (**A**) and *imp* ΔPLD (**B**) whole-mount fillets at 24 h post-eclosion. CCAP/Bursicon axons were detected using CCAP-Gal4 driven mCD8-RFP expression (red in **A-B**) and GFP-Imp signal is shown in green in **A-B** and **A’-B’**. Note that the overall reduction of GFP-Imp levels throughout the axonal tract in *imp* ΔPLD is due to the constitutive nature of the ΔPLD mutation. Scale bar: 5 µm. Animals were raised at 29°C. Complete genotypes: control: GFP-Imp/Y; CCAP-Gal4,UAS-mCD8-RFP/CyO; *imp* ΔPLD: GFP-ImpΔPLD/Y; CCAP-Gal4,UAS-mCD8-RFP/Sco. **C** Dot plot showing GFP-Imp mean intensity in CCAP/Bursicon distal axons of adult flies at 24 h post-eclosion under control and *imp* ΔPLD conditions. Number of CCAP/Bursicon distal axons analyzed: control=24; *imp* ΔPLD=29. 3 replicates were performed. Individual data points correspond to the mean value obtained from 1 to 2 analyzed distal axon portions per individual and were normalized to the control condition. Bars and error bars correspond respectively to mean and SEM. **P<0.01 (two-tailed Mann-Whitney U test on individual data points). **D-E** Tile confocal images of control (**D**) and *imp* ΔPLD (**E**) whole-mount fillets dissected at 24 h post-eclosion and stained using anti-Bursicon antibody. Red asterisks indicate the axon terminals. Scale bar: 100 µm. **F** Dot plot showing the maximal axonal length of control and *imp* ΔPLD CCAP/Bursicon neurons at 24 h post-eclosion. Number of fillets analyzed: control=27 and *imp* ΔPLD=42. **G** Dot plot showing the number of CCAP/Bursicon axon terminals in control and *imp* ΔPLD adult flies at 24 h post-eclosion. Number of fillets analyzed: control=22 and *imp* ΔPLD=38. **D-E** 4 replicates were performed. Quantifications were performed on the same individuals when possible. Bars and error bars correspond respectively to mean and SEM. **P<0.01(two-tailed Mann-Whitney U test on individual data points).

Quantitative analysis of adult CCAP/Bursicon axonal morphology revealed that, while axonal length was unaffected in *imp* ΔPLD mutants, the number of axon terminals was significantly reduced compared to controls (Figure 5D-G). These branching defects may result from reduced axonal localization of Imp, the moderate reduction in overall Imp levels, or a combination of both.

Overall, these results demonstrate that Imp-dependent regulation of axonal branching and late axonal regrowth can be genetically uncoupled.

### *profilin* overexpression restores late axonal regrowth but not branching defects

The genetic uncoupling of axonal branching and late regrowth suggested that these two Imp-dependent processes may rely on distinct downstream effectors. *profilin* mRNA, which encodes an actin binding protein promoting F-actin filament polymerization, was previously identified as a molecular and functional target of Imp (Cooley et al. 1992; Medioni et al. 2014). To determine whether Imp mediates the remodeling of CCAP/Bursicon neurons through *profilin* regulation, we performed epistasis experiments by overexpressing *profilin* in the *imp* Degrad loss-of-function background. Because *imp* inactivation induces severe axonal morphology defects (see Figure 1), we asked whether increasing Profilin expression could rescue these phenotypes.

Quantitative analysis revealed that *profilin* overexpression restored the axonal length defects of *imp* Degrad individuals to levels comparable to control flies (Figure 6A,C,D). In contrast, axonal arborization remained impaired, adult axons displaying a reduced number of axon terminals (Figure 6A-C,E), suggesting that *profilin* is a downstream effector of Imp in late axonal regrowth, but not branching. Consistent with this, *profilin* overexpression also failed to rescue the branching defects observed in *imp* ΔPLD mutants (Figure S4).

**Figure 6:**
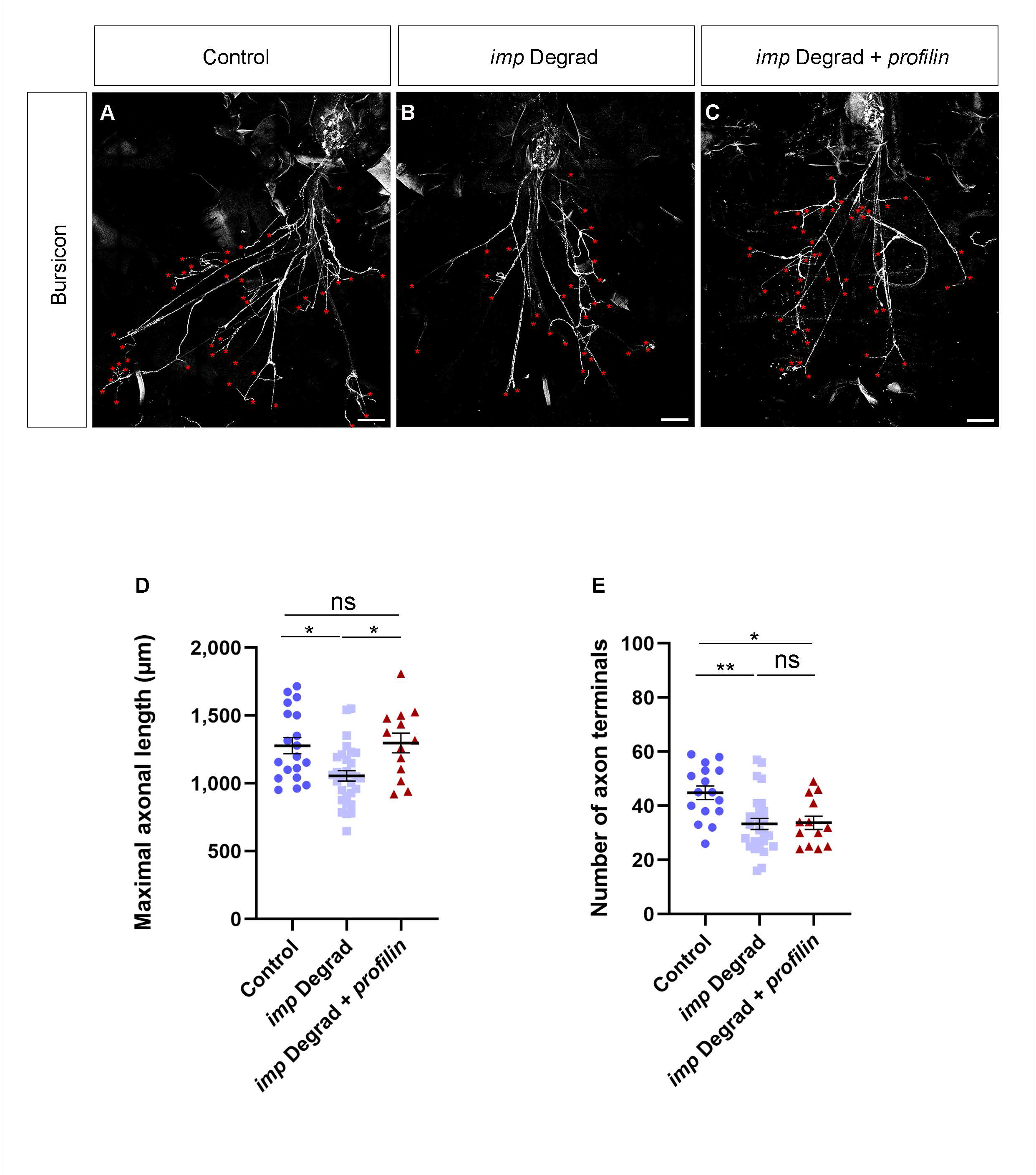
*profillin* overexpression rescues *imp* Degrad axonal regrowth defects but not branching of CCAP/Bursicon neurons. **A-C** Tile confocal images of control (**A**), *imp* Degrad (**B**) and *imp* Degrad+*profilin* (**C**) obtained from whole-mount fillets at 24 h post-eclosion and stained using anti-Bursicon antibody. Red asterisks indicate the axon terminals. Scale bar: 100 µm. Animals were raised at 29°C. Complete genotypes: control: GFP-Imp/Y; CCAP-Gal4,UAS-mCD8-RFP/+; *imp* Degrad: GFP-Imp/Y; CCAP-Gal4,UAS-mCD8-RFP/+; UAS-*degrad-fp*,UAS-*gfp*-RNAi/+; *imp* Degrad+*profilin*: GFP-Imp/Y; CCAP-Gal4/+; UAS-*degrad-fp*,UAS-*gfp*-RNAi/UAS-*profilin*. **D** Dot plot showing the maximal axonal length of control, *imp* Degrad and *imp* Degrad+*profilin* CCAP/Bursicon neurons at 24 h post-eclosion. Number of fillets analyzed: control=19; *imp* Degrad=30; *imp* Degrad+*profilin*=13. **E** Dot plot showing the number of CCAP/Bursicon axon terminals in control, *imp* Degrad and *imp* Degrad+*prof*ilin adult flies at 24 h post-eclosion. Number of fillets analyzed: control=16; *imp* Degrad=27; *imp* Degrad+*profilin*=13. **D-E** 2-3 replicates were performed. Quantifications were performed on the same individuals when possible. Bars and error bars correspond respectively to mean and SEM. **P<0.01; *P<0.05; ns stands for not significant (Kruskall– Wallis with Dunn’s post-tests).

Our observations thus support the idea that these morphological processes are regulated through distinct molecular mechanisms, with *profilin* mRNA preferentially contributing to axonal regrowth.

### Imp controls *profilin* mRNA stability in CCAP/Bursicon neurons

To investigate how Imp regulates *profilin* mRNA to promote axonal regrowth, we examined *profilin* mRNA abundance using single-molecule fluorescence in situ hybridization (smFISH) combined with high-resolution imaging.

In control CCAP/Bursicon cell bodies, *profilin* transcription foci were detected as bright nuclear spots (Fig 7A, arrowhead) and processed mRNA molecules as cytoplasmic fluorescent spots (Figure 7A) whose number decreased upon RNAi-mediated *profilin* inactivation (Figure S5A-C).

**Figure 7:**
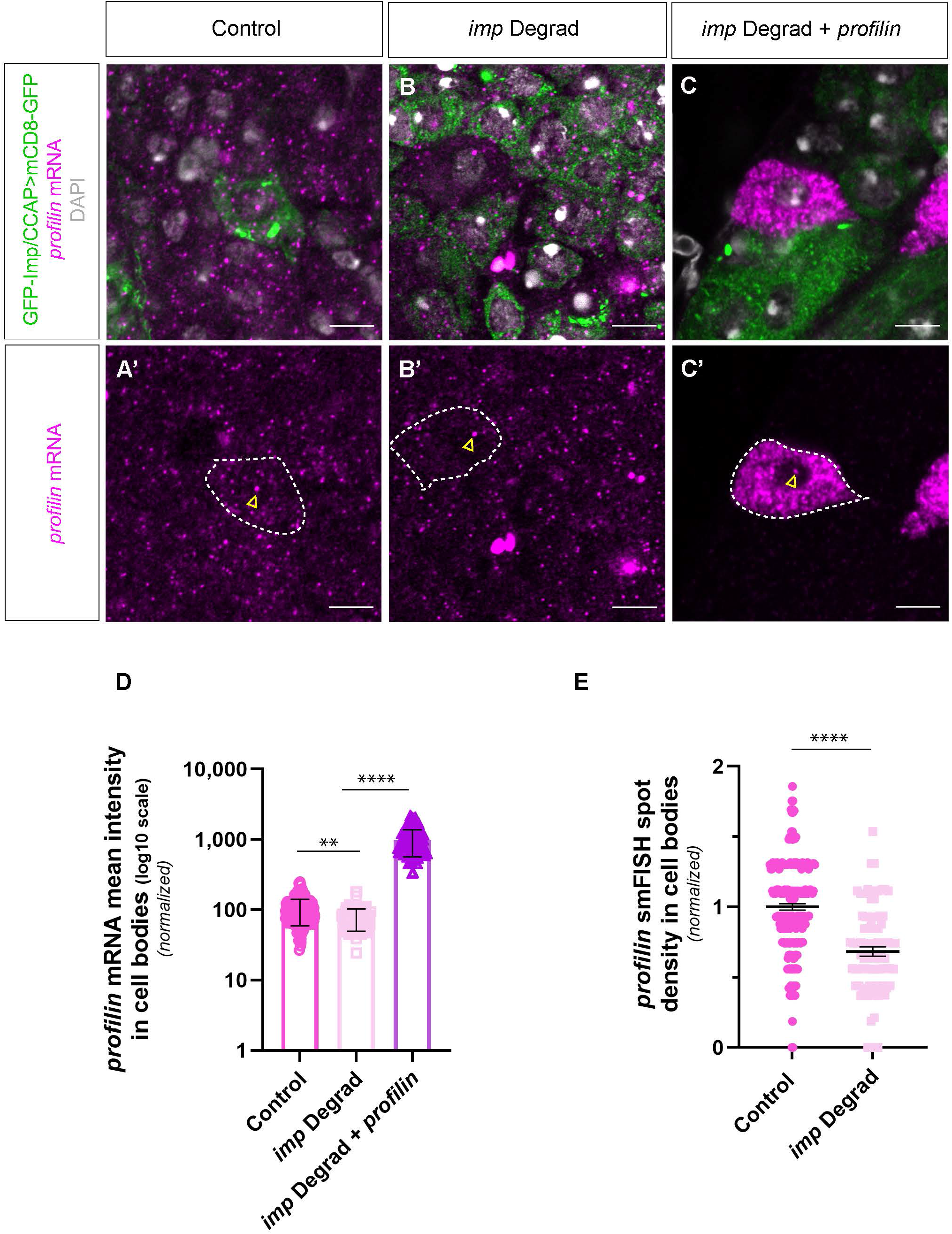
Imp regulates *profilin* mRNA levels in CCAP/Bursicon neurons. **A-C** Confocal images of CCAP/Bursicon neuron cell bodies from control (**A**), *imp* Degrad (**B**) and *imp* Degrad+*profilin* (**C**) whole-mount fillets incubated with anti-*profilin* probes. In **A**, CCAP/Bursicon neurons were detected using CCAP-Gal4 driven mCD8-GFP expression (in green). In **B,C**, CCAP/Bursicon cell bodies were identified based on DAPI staining (in white) as well as the absence of GFP signal in *imp* Degrad and *imp* Degrad+*profilin* conditions. *profilin* smFISH signals are shown in magenta in **A-C** and **A’-C’**. Brightness and contrast were adjusted for each image for display purposes. Yellow arrowheads point to transcription foci. Scale bar: 5 µm. Animals were raised at 29°C. Complete genotypes: control: GFP-Imp/Y; CCAP-Gal4,UAS-mCD8-GFP/+; *imp* Degrad: GFP-Imp/Y; CCAP-Gal4/+; UAS-*degrad-fp*,UAS- *gfp*-RNAi/+; *imp* Degrad+*profilin*: GFP-Imp/Y; CCAP-Gal4/+; UAS-*degrad-fp*,UAS-*gfp*-RNAi /UAS-*profilin*. **D** Dot plot showing *profilin* mRNA signal mean intensity in CCAP/Bursicon cell bodies of control, *imp* Degrad and *imp* Degrad+*profilin* adult flies at 24 h post-eclosion. Number of CCAP/Bursicon cell body regions analyzed: control=197; *imp* Degrad=104; *imp* Degrad+*profilin*=150. 4 replicates were performed. Individual data points correspond to individual regions (up to 2 per cell body) and were normalized to the control condition. 1 to 6 CCAP/Bursicon cell bodies were analyzed per individual. Bars and error bars correspond respectively to mean and SEM. ****P<0.0001; **P<0.001 (Kruskall– Wallis with Dunn’s post-tests). **E** Dot plot showing *profilin* smFISH spot density in CCAP/Bursicon cell bodies of control and *imp* Degrad flies at 24 h post-eclosion. Number of cell body regions analyzed: control=231; *imp* Degrad=88. 4 replicates were performed. Individual data points correspond to individual regions analyzed (up to 2 per cell body). Data were normalized to the control condition. 1 to 5 CCAP cell bodies were analyzed per individual. Bars and error bars correspond respectively to mean and SEM. ****P<0.0001 (two-tailed Mann-Whitney U test on individual data points).

In *imp* Degrad condition, a specific and significant reduction of cytoplasmic smFISH signal was observed compared to controls, characterized by a decrease in both *profilin* smFISH spot density and mean signal intensity (Figure 7A,B,D,E), indicating that Imp is required to maintain normal *profilin* mRNA levels. *profilin* overexpression in *imp* Degrad condition counteracted this effect and led to a strong increase of cytoplasmic *profilin* mRNA signal intensity (Figure 7C,D), indicating that the rescue of late axonal regrowth observed in this context is associated with the recovery of *profilin* mRNA levels. These observations support the idea that impaired *profilin* mRNA levels cause the late regrowth defects induced upon strong *imp* inactivation.

Further consistent with this model, *profilin* mRNA abundance was unaffected in the CCAP/Bursicon cell bodies of *imp* ΔPLD flies, in which late axonal regrowth is preserved (Figure S5A–D).

Together, the decrease in cytoplasmic but not nuclear *profilin* mRNA signal in *imp* Degrad condition suggests that *profilin* mRNA levels are post-transcriptionally regulated by Imp function in CCAP/Bursicon cell bodies. Such regulation is selectively contributing to late axonal regrowth, indicating the existence of additional mechanisms controlling axonal branching.

## Discussion

In this study, we identify the RNA-binding protein Imp as a central regulator of the developmental axonal regrowth program triggered in Drosophila CCAP/Bursicon neurons after pruning of larval axonal arborization. Our data reveal three principal findings. First, Imp acts during a restricted developmental window during late metamorphosis to promote the establishment of a functional adult axonal network. Second, Imp controls two distinct processes, late axonal regrowth and axonal branching, that can be genetically uncoupled. Third, Imp-mediated regulation of *profilin* mRNA stability seems essential to axonal elongation but dispensable for the branching process. Together, our results suggest that Imp coordinates temporally separated axonal morphogenetic programs by regulating separate pools of target mRNAs.

### Imp functions during a restricted developmental window to promote axonal regrowth after pruning

During their lifetime, neurons undergo different phases of axonal growth – primary growth, developmental regrowth or post-traumatic regeneration – each governed by dedicated molecular machineries. Members of the IGF2BP family of RNA-binding proteins have been shown to regulate multiple aspects of axonal growth programs. In vertebrates, ZBP1 was shown to direct initial axon elongation in response to guiding cues *in vitro* (Zhang et al. 2001; Leung et al. 2006; Yao et al. 2006; Welshhans and Bassell 2011) and to promote axonal navigation during *in vivo* early development (Hornberg and Holt 2013; Gaynes et al. 2015; Lepelletier et al. 2017). During maturation of the nervous system, expression of ZBP1 declines and this reduction was associated with the loss of regenerative capacity in adult mammalian neurons (Donnelly et al. 2011; Jones et al. 2021). Indeed, transient upregulation of ZBP1 levels in dorsal root ganglion neurons could restore axon regeneration after axotomy. In *Drosophila*, we previously showed that Imp is transported to axons at metamorphosis and promotes the regrowth of Mushroom Body γ neuron axonal branches that occurs after developmentally controlled pruning of early-born projections (Medioni et al. 2014).

Our present study extends these observations by demonstrating that Imp activity is required in another neuronal population, the CCAP/Bursicon neurons, where it promotes the establishment of a mature axonal network in a specific time-window at the end of metamorphosis. Importantly, this temporal requirement coincides with a developmental increase in Imp protein abundance.

More generally, our findings support the emerging concept that temporal control of RNA-binding protein expression and regulation represents an intrinsic mechanism regulating initial growth, developmental regrowth and regenerative capacity (Holt et al. 2019; Parra and Johnston 2022).

### Axonal regrowth and branching are regulated by distinct remodeling programs

Although axonal elongation and branching are often described as processes whose coordination ensures proper axonal arborization (Holt et al. 2019; Parra and Johnston 2022), our results demonstrate that they can be temporally dissociated in the context of *in vivo* axonal regrowth. Indeed, branching of CCAP/Bursicon axons occurs predominantly during the early phase of axonal regrowth, and is almost completed before the late phase of regrowth takes place later. Moreover, these two processes respond differentially to perturbation of Imp function. While strong *imp* inhibition generated by *imp* Degrad expression impairs both axonal branching and late regrowth, the milder downregulation of *imp* level and alteration of Imp axonal transport induced by the constitutive *imp* ΔPLD mutation specifically compromise branching.

Such an uncoupling is consistent with previous work demonstrating that transcriptional programs controlling different steps of axonal remodeling are temporally segregated and that branching and elongation rely on distinct molecular machineries (Kalil and Dent 2014; Alyagor et al. 2018). Our discovery that *profilin* overexpression efficiently rescues the axonal elongation defects caused by *imp* inhibition, but fails to restore branching, further strengthens this notion and suggests that Imp govern these distinct events through regulation of separate transcripts.

A similar molecular decoupling has been proposed in mammalian neurons, where ZBP1-dependent local translation of axonally-localized *β−actin* mRNA preferentially supports branch formation, whereas local translation of axonal *gap-43* mRNA promotes elongating growth (Donnelly et al. 2013). By analogy, our findings suggest that Imp coordinates multiple remodeling programs by regulating distinct mRNA molecules, which may represent a conserved strategy by which post-transcriptional regulation enables spatial and temporal coordination.

Cytoplasmic RBPs regulate their targets at different levels: controlling their stability, subcellular localization or translation. In neurons, some were shown to promote the transport of target mRNAs to axons, as well as to control their local translation. Such a process has emerged as a fundamental mechanism controlling neuronal growth, plasticity and regeneration by allowing rapid and spatially restricted proteome remodeling within neuronal processes (Holt et al. 2019). In the context of branching, mRNA molecules were shown in vertebrate retinal ganglion cells to transiently dock at discrete subcellular sites that will define future axonal branch positions, and to undergo local translation prior to branch initiation (Kalous et al. 2014; Shigeoka et al. 2016; Wong et al. 2017; Cioni et al. 2019). These molecules subsequently localize to stabilized branches, suggesting that mRNA localization coupled to local translation supports both branch formation and maturation. Thus, spatially restricted translation can provide a molecular trigger required to precisely position individual branches while avoiding inappropriate activation of growth programs throughout the axonal arbor.

The branching phenotypes we observed in the *imp* ΔPLD mutant suggest that axonally localized Imp may promote branching through the local regulation of specific transcripts distinct from *profilin* mRNA. Although we cannot exclude that the moderate reduction of Imp levels in cell bodies observed in the *imp* ΔPLD mutant contributes to the phenotype, the specific branching defects observed suggests that branch formation may be particularly sensitive to local post-transcriptional regulation. Future identification of axonally-localized Imp target mRNAs will be essential to determine if and how local translation programs generate branch-specific morphogenesis.

More broadly, our work identifies post-transcriptional regulation as a mechanism capable of uncoupling axonal elongation and branching processes and highlights mRNA regulation as a major contributor to neuronal remodeling. The mechanisms we describe have been evolutionary conserved, highlighting that members of the IGF2BP family contribute to neuronal remodeling and regenerative responses in various contexts, among which axon regeneration and local translation following injury (Donnelly et al. 2011; Turner-Bridger et al. 2020; Parra and Johnston 2022).

## Materials and methods

### Drosophila melanogaster stocks and genetics

Fly crosses were performed on standard media and raised either at 18°C to minimize Gal4/UAS system activity for experiments shown in Figure 2, 25°C to maintain physiological developmental temperature for experiments shown in Figure 2, 3, 4, S2, S3 A-B and A’-B’ or 29°C for experiments shown in Figure 1, 5, 6, 7, S1, S3 C-F and C’-E’, S4 and S5. As *imp* is X-linked, males were exclusively used for experiments.

The following fly stocks were used: GFP-Imp protein trap-line #G080 (Medioni et al. 2014), yw; CCAP-Gal4/CyO (BDSC#25685), UAS-NSlmb-vhh GFP4, UAS-*gfp*-RNAi/TM6b #Degrad (gift from Markus Affolter), UAS-mCD8-RFP (BDSC#27391), *w^1118^* (BDSC#6326), UAS-*profilin* (Cooley et al. 1992), UASp-*flag*-*imp* (Medioni et al. 2014), UAS-mCD8-GFP (BDSC#5137), GFP-ImpΔPLD (Vijayakumar et al. 2019), UAS-*profilin*-RNAi (VDRC#KK102759).

### Temperature-shift experiments

GAL4/UAS-dependent transgene expression was modulated by temperature shifts as previously described (Nagarkar-Jaiswal et al. 2015), using 18°C as the restrictive temperature and 25°C as the activating temperature for GAL4/UAS activity. Shifts were performed between 18°C and 25°C at the indicated hours After Puparium Formation (APF). At 18°C, embryonic, larval, and pupal stages last approximately 2–3, 6–7, and 8–9 days, respectively, compared with ∼1, 4, and 5 days at 25°C.

Based on these developmental timings, two days at 18°C correspond approximately to one day at 25°C during the pupal stage. Developmental stages were verified using the morphological criteria described by (Bainbridge and Bownes 1981).

### Wing phenotype scoring

Wing phenotypes were scored 24 h after eclosion as described previously (Luan et al. 2006). Wings were considered expanded only when fully flattened.

### Pupal staging

Crosses were reared at 25°C and white prepupae were collected in water-containing dishes. Pupae were aged at 25°C until indicated hours after puparium formation before dissection.

### Fillet preparations

Fillets were dissected in cold 1x PBS on silicone-coated dissection plates (SYLG184, WPI) for a maximum of 1 h. The pupal case was kept for pupal stage dissections whereas legs and wings were removed for adult dissections. Individuals were pinned rostrally and posteriorly using fine dissecting needles. Pupal cases and cuticles were incised using precision scissors along the midline, from the posterior part of the abdomen to the thorax to expose internal structures. Following the midline incision, the cuticle flaps were pinned to the silicone-coated dissection plate using fine dissecting needles, allowing the preparation to remain fully open. Surrounding tissues were then gently removed by pipetting and with forceps to expose the Ventral Nerve Cord (VNC) and the fillets.

When no immuno-staining was performed, individuals were fixed directly on dissection plates with 4% formaldehyde (FA) 0.1% PBS-Triton-X-100 (PBT) for 45 min and washed thrice for 15 min in 0.1% PBT at room temperature under gentle agitation. Samples were mounted in Vectashield (Vector Laboratories) medium containing DAPI.

### Immunostaining

Adult fillets of 1-day-old *Drosophila* were pre-fixed directly on dissection plates in 4% FA 1x PBS for 25 min and subsequently post-fixed in 4% FA 0.5% PBT for 25 min, then washed in 0.1% PBT for 15 min. Samples were transferred into 1.5 mL tubes and incubated for 2 h in blocking solution (0.1% PBT supplemented with 0.5% BSA and 5% goat serum). Fillets were then incubated with primary antibodies (rabbit anti-Bursicon (homemade, 1:1,000)) overnight at 4°C under agitation. On the next day, samples were washed once in 0.1% PBT, then 5 times in blocking solution for 30 min each. Fillets were then incubated for 2 h in blocking solution supplemented in anti-rabbit antibodies conjugated with Alexa Fluor 633 or 568 (Thermo Fisher, 1:1,000 both). Dissected fillets were washed four times in PBT 0.1% for 20 min and mounted in Vectashield (Vector Laboratories) medium optionally containing DAPI.

### Production of anti-Bursicon antibody

Affinity-purified rabbit anti-Bursicon antibody was generated using the peptide described by (Luan et al. 2006) (CEGPLNNHFRRIALQ), that corresponds to the C-terminal region of the Bursicon α-subunit. The peptide was conjugated to bovine serum albumin (BSA) *via* maleimide coupling and used as the immunogen.

### Single-molecule Fluorescent In Situ Hybridization (smFISH)

Adult fillets from 1-day-old *Drosophila* were dissected in cold RNase-free HL3 buffer (NaCl 70 mM, KCl 5 mM, MgCl_2_ 4 mM, trehalose 5 mM, sucrose 115 mM, HEPES 5 mM, NaHCO_3_ 10 mM, pH 7.2–7.3). Dissected fillets were then fixed in 4% FA in 1x PBS for 1 h at 4°C, rinsed twice with 1x PBS directly on dissection plates. Dissected fillets were carefully transferred into 1.5 mL low-binding tubes and stored overnight at 4°C in 70% ethanol diluted in 1x PBS. On the next day, fillets were washed twice with 1x PBS 15 min, followed by wash buffer (10% formamide in 2x SSC) for 5 min. Fillets were then incubated overnight, at 45°C, and under agitation, with Quasar®670-labeled Stellaris® probes in 100 μL hybridization buffer (100 mg/mL dextran sulfate, 10% formamide in 2x SSC). *profilin* probes were used at a final concentration of 0.125 μM.

Sequences of *profilin* probes used (described in (Formicola et al. 2021)) from 5’ to 3’ were as follows: *ccgcaacaccgacgattt*, *cacacgaaattggcaggg*, *tcgcactttcgtttcggg*, *ttgctttaccgcacggcg*, *gatctggat atggatcgc*, *gggtgcggattaagttga*, *catggtgctttgtttgtc*, *gtccacataatcttgcca*, *ctgcgaggccaggagttg*, *gatgca cgccttggtcac*, *ccaaatgttgccgtcgtg*, *tcacctcaaagccactgg*, *gtttggagagctcctctt*, *ctggtcaaagccgctgat*, *gttgc tggtgagaccgtc*, *aaatgtaccgctggccgg*, *gcggtctgtgccggaaag*, *ttcatgcagtgcactccg*, *acgatcacggcttgtgtt*, *c gggatcctcgtagatgg*, *tctctaccacggaagcgg*, *ctattctcctagtacccg*, *tcatttacggttcgctct*, *tggtttttcttttcccat*, *gc aaattctttcttggcc*, *tcctctgctacacacaaa*, *gcatttttactcgatcca*.

After hybridization, fillets were incubated 30 min at 45°C in pre-warmed wash buffer under agitation, washed 5 min at room temperature in 2x SSC and mounted in Vectashield (Vector Laboratories) medium containing DAPI.

### Image acquisition

Fillets were imaged in tile scan mode using a Zeiss LSM780 NLO confocal microscope equipped with GaAsP detectors and a 25 x 0.8 NA oil immersion objective. Images were taken with a 0.47 µm xy pixel size and an optimal z step size of around 1.15 µm. Tile scans were stitched and maximum intensity projections of z-stacks were generated using Zen (black edition, 2011, SP7 FP3 64bit V14.0.19.201) software.

Ventral nerve cords were imaged using Zeiss LSM880 confocal microscope equipped with an Airyscan module and a 40 × 1.4 NA oil immersion objective. Images were taken with a 0.231 μm xy pixel size and a 1 μm z step size and processed with the automatic Airyscan processing module of Zen (black edition, 2.3 SP1 FP3 64bit V14.0.26.201) software.

For imaging of smFISH spots and GFP-Imp signal, samples were imaged using a Zeiss LSM880 confocal microscope equipped with an Airyscan module and a 63 × 1.4 NA oil immersion objective. Images were taken with a 0.04 μm xy pixel size and a 0.5 μm z step size and processed with the automatic Airyscan processing module of Zen Black software.

### Image analysis

#### Maximal axonal length and number of axonal terminal quantification

Image analysis was performed on maximum intensity projections (MIPs). When appropriate, MIPs were generated from a subset of raw z-sections containing the structures of interest only. Brightness and contrast display settings were adjusted to facilitate visualization during image analysis. Axonal length was measured using the freehand line tool of Fiji by tracing the longest CCAP/Bursicon-positive axonal process from the point where it emerged from the GFP-Imp-labelled VNC to its most distal terminal (Figure S1C-D). Axonal branching was quantified by counting the number of axon terminals using the multipoint tool of Fiji (Figure S1E).

Representative images of fillets shown in Figures 1, 3, 5, 6, S1 and S4 were processed separately in Fiji for display. Brightness and contrast were adjusted for optimal visualization. In addition, smoothing and background subtraction were applied only when necessary for illustration and were not used for morphometric analyses.

#### Size and number quantifications of cell bodies

The area of individual CCAP/Bursicon cell bodies was measured manually by outlining cell bodies on the optical section displaying the largest cross-sectional area within confocal z-stacks using the freehand selection tool of Fiji. The number of CCAP/Bursicon cell body number per ventral nerve cord was determined manually from confocal z-stack using the multi-point tool of Fiji.

#### Imp level quantification in cell bodies

One ROI of 100 × 100 pixels was cropped from the cytoplasm of CCAP/Bursicon cell bodies on single z-slices. The GFP mean intensity was calculated for each ROI.

#### Imp level quantification in axons

Images with a pixel size of 0.04 µm containing axon volume ranging from ∼60 to 1,600 µm^3^ were analyzed. 3D surface reconstruction of mCD8-RFP-labelled CCAP/Bursicon axons was generated from Z-stacks using the surface tool of Imaris software with automatic thresholding and surface detail of 0.146 μm. Axonal regions immediately adjacent to tissues with high GFP-Imp signal were excluded from the analysis. All remaining axonal segments were merged into a single surface using the unify tool of Imaris. The absolute mean GFP signal intensity of the reconstructed surface was extracted.

#### 2D particle detection

Two ROIs of 100 × 100 pixels were cropped from the cytoplasm of CCAP/Bursicon cell bodies on single z-slices for smFISH signal quantification. The ROIs used for GFP-Imp mean intensity quantification were also used for GFP-Imp granule detection. ROIs were resized to a factor 1 using the Laplacian Pyramid plugin in ImageJ, then rescaled to enhance contrast and keep 0.01% pixels saturated (for smFISH quantifications only), and finally converted from 32-bit to 16-bit images to change float numbers to integer values.

GFP-Imp granules and *profilin* smFISH spots were detected using the Small Particle Detection (SPaDe) algorithm (https://gitlab.inria.fr/ncedilni/spade). The cutoff size for particles was set to 4 pixels. Detection thresholds were set to 0.32 for GFP-Imp granules and 0.62 for *profilin* smFISH spots. For each analyzed field, particle masks were extracted and the number of detected objects was quantified.

## Acknowledgments

This study was supported by grants (Credit Scientifique Incitatif Cote d’Azur University and Initiative of Excellence Université Côte d’Azur under reference number ANR-15-IDEX-01) to C. M. and grants from the ANR (# ANR-20-CE16-0010 and ANR22-CE12-0024), and from the CEFIPRA (IFC/6503-E/2021/193) to F.B., by fellowships (ministerial doctoral contract, MENRT and ATER, Cote d’Azur University) to M.N. We are grateful to M. Affolter, L. Cooley J.M. Dura, R. Hewes, B. White, DHSB (Iowa University), the Bloomington Drosophila Stock Center and Vienna Drosophila Resource Center for reagents. We thank the iBV PRISM Imaging facility for use of their microscopes and support. We are also deeply thankful for T. Gu advices to dissect CCAP/Bursicon fillets. We finally thank F. De Graeve, J.M Dura, B. Hudry and L. NGuyen for critical reading of the manuscript.

---

**Supplementary Figure 1:**
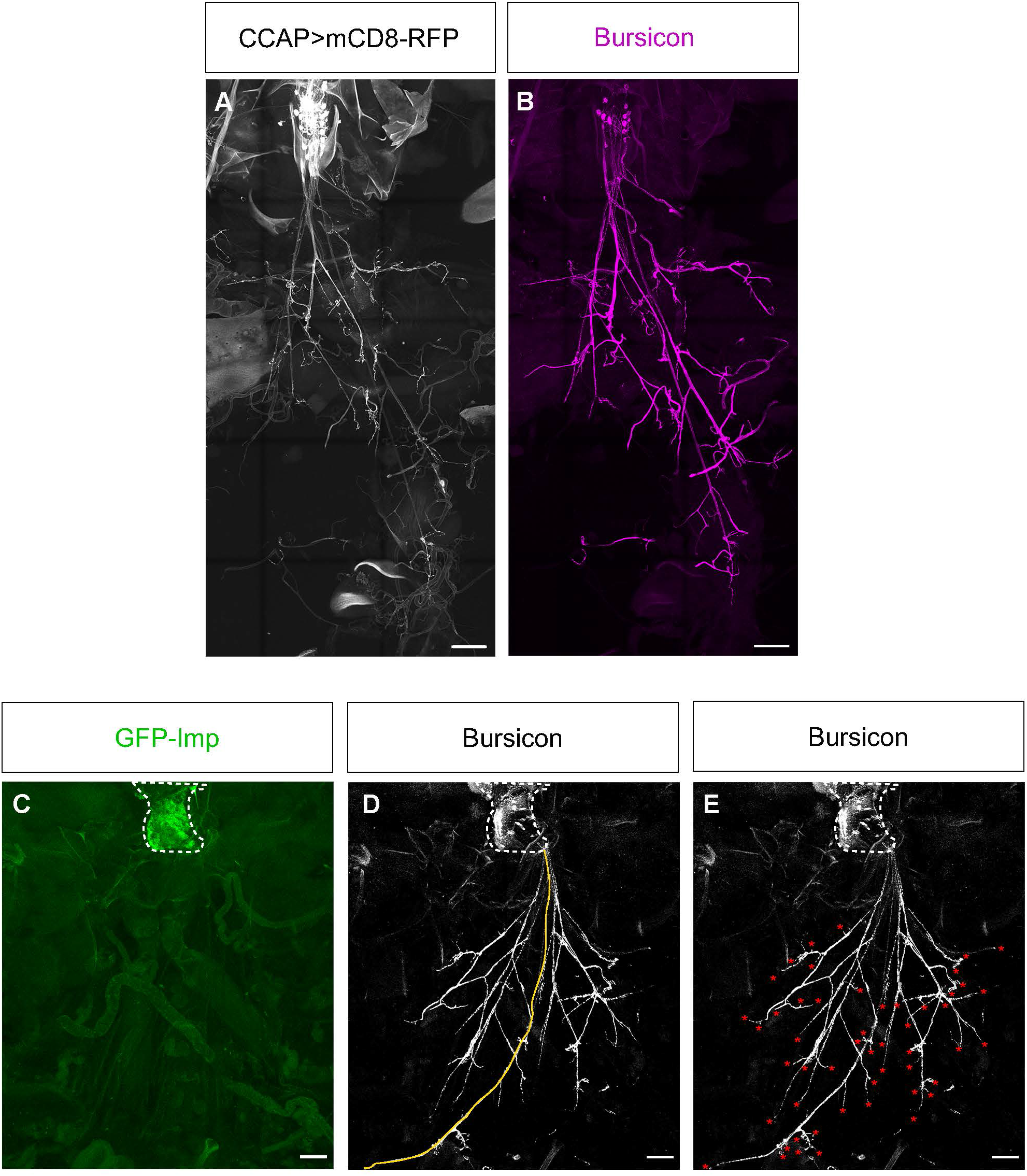
Specific labeling of CCAP/Bursicon neurons and methodology for axonal morphology analysis. **A-B** Tile confocal images of adult CCAP/Bursicon neurons labeled using the CCAP-Gal4 driver in combination with the membrane-associated mCD8-RFP (**A**) and anti-Bursicon antibody (**B**). Scale bar: 100 µm. Complete genotype: control: GFP-Imp/Y; CCAP-Gal4,UAS-mCD8-RFP/+. Similar labeling of CCAP/Bursicon axons is observed when imaging CCAP>mCD8-RFP and Bursicon signal. **C-E** Illustration of the quantification method used to assess maximal axonal length (**D**) and number of axon terminals (**E**). The outer boundary of the Ventral Nerve Cord (VNC) (delimited with a white dotted line in **C**) is detected based on bright GFP-Imp labeling (green in **C**). It was used as a reference starting point to measure the length of the longest CCAP/Bursicon axonal process, highlighted in yellow in **D**. **E** The number of axon terminals was quantified by counting axonal terminal ends, indicated by red asterisks. Complete genotype: GFP-Imp/Y; CCAP-Gal4,UAS-mCD8-RFP/+. Scale bar: 100 µm.

**Supplementary Figure 2:**
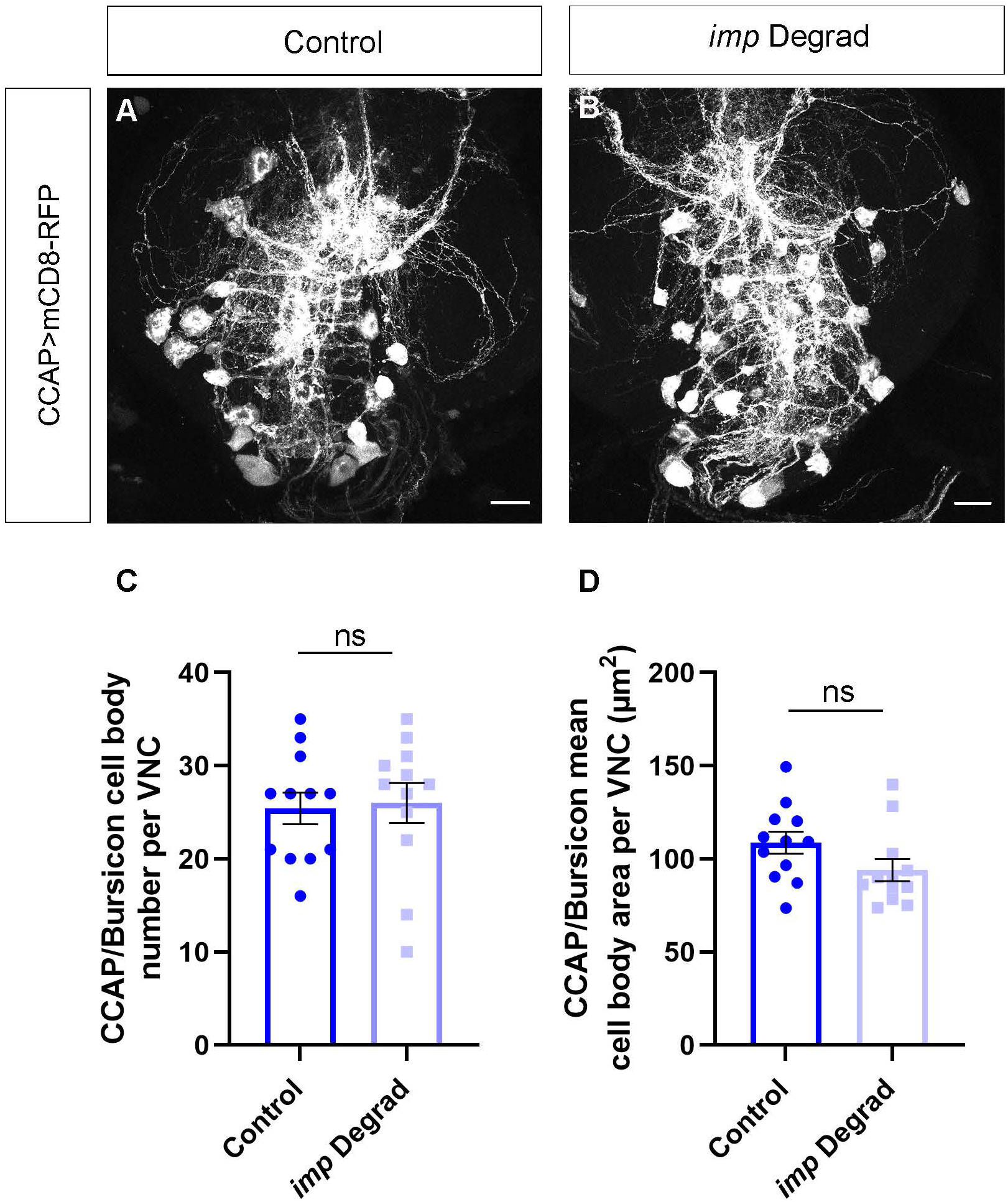
CCAP/Bursicon cell body differentiation is preserved following *imp* inactivation. **A-B** Confocal images of control (**A**) and *imp* Degrad (**B**) whole-mount ventral nerve cords dissected at 24 h post-eclosion. Scale bar: 20 µm. Complete genotypes: control: GFP-Imp/Y; CCAP-Gal4,UAS-mCD8-RFP/CyO; *imp* Degrad: GFP-Imp/Y; CCAP-Gal4,UAS-mCD8-RFP/+; UAS-*degrad-fp*,UAS-*gfp*-RNAi/+. Animals were raised at 25°C. **C** Dot plot showing the number of CCAP/Bursicon cell bodies per VNC in control and *imp* Degrad conditions. **D** Dot plot showing the mean area of CCAP/Bursicon cell bodies per VNC in control and *imp* Degrad conditions. **C-D** Number of VNC analyzed: control=12; *imp* Degrad=12. 1 replicate was performed. Individual data points correspond to individual VNC. Bars and error bars correspond respectively to mean and SEM. Quantifications were performed on the same individuals. ns stands for not significant (Two-tailed Mann-Whitney U test on individual data points).

**Supplementary Figure 3:**
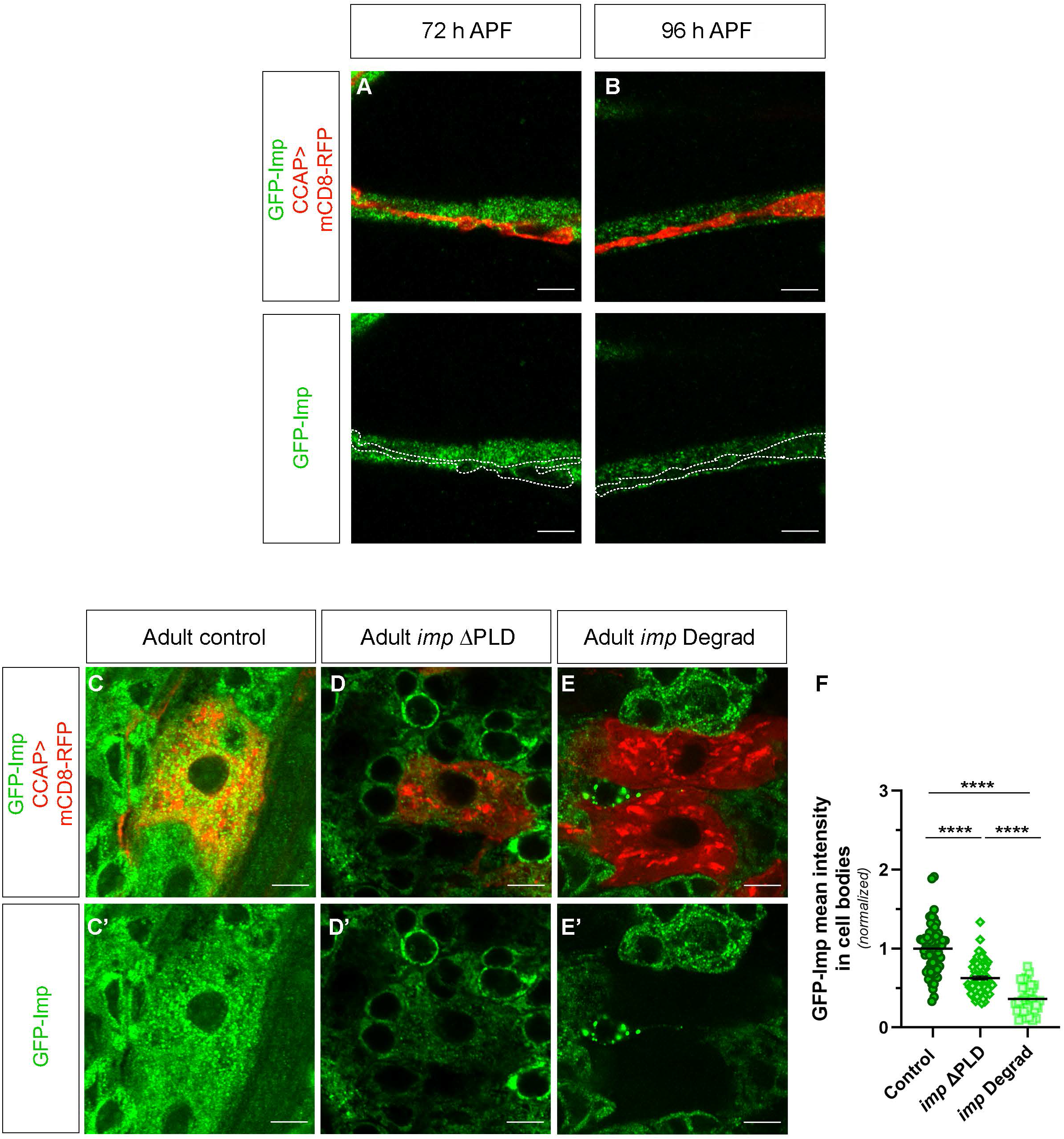
GFP-Imp localizes to granules within distal axons and is reduced in CCAP/Bursicon cell bodies in *imp* ΔPLD flies. **A-D** Confocal images of control (**A-B**) CCAP/Bursicon neuron distal axons from whole-mount fillets dissected at 72 h APF (**A**) and 96 h APF (**B**). CCAP/Bursicon neurons were detected using CCAP-Gal4 driven mCD8-RFP expression (in red in **A-B**), and GFP-Imp signal is shown in green in **A-B** and **A’-B’**. Scale bar: 5 µm. Animals were raised at 25°C. Complete genotype: GFP-Imp/Y; CCAP-Gal4,UAS-mCD8-RFP/CyO. **C-E** Confocal images of CCAP/Bursicon neuron cell bodies obtained from control (**C**), *imp* ΔPLD (**D**) and *imp* Degrad (**E**) whole mount fillets at 24 h post-eclosion. CCAP/Bursicon neurons were detected using CCAP-Gal4 driven mCD8-RFP expression (in red in **C-E**), and GFP-Imp signal is shown in green in **C-E** and **C’-E’**. Scale bar: 5 µm. Animals were raised at 29°C. Complete genotypes: control: GFP-Imp/Y; CCAP-Gal4,UAS-mCD8-RFP/+; *imp* ΔPLD: GFP-Imp ΔPLD/Y; CCAP-Gal4,UAS-mCD8-RFP/Sco; *imp* Degrad: GFP-Imp/Y; CCAP-Gal4,UAS-mCD8-RFP/+; UAS-*degrad-fp*,UAS-*gfp*-RNAi/+; **F** Dot plot showing the GFP-Imp mean intensity in CCAP/Bursicon cell bodies of adult flies at 24 h post-eclosion under control, *imp* ΔPLD and *imp* Degrad conditions. Number of CCAP/Bursicon cell body analyzed: control=66; *imp* ΔPLD=76; *imp* Degrad=36. 3 replicates were performed. Individual data points correspond to individual cell bodies for each condition and were normalized to the control condition. 1 to 6 CCAP/Bursicon cell bodies were analyzed per individual. Bars and error bars correspond respectively to mean and SEM. ****P<0.0001 (Kruskal-Wallis with Dunn’s post-tests).

**Supplementary Figure 4:**
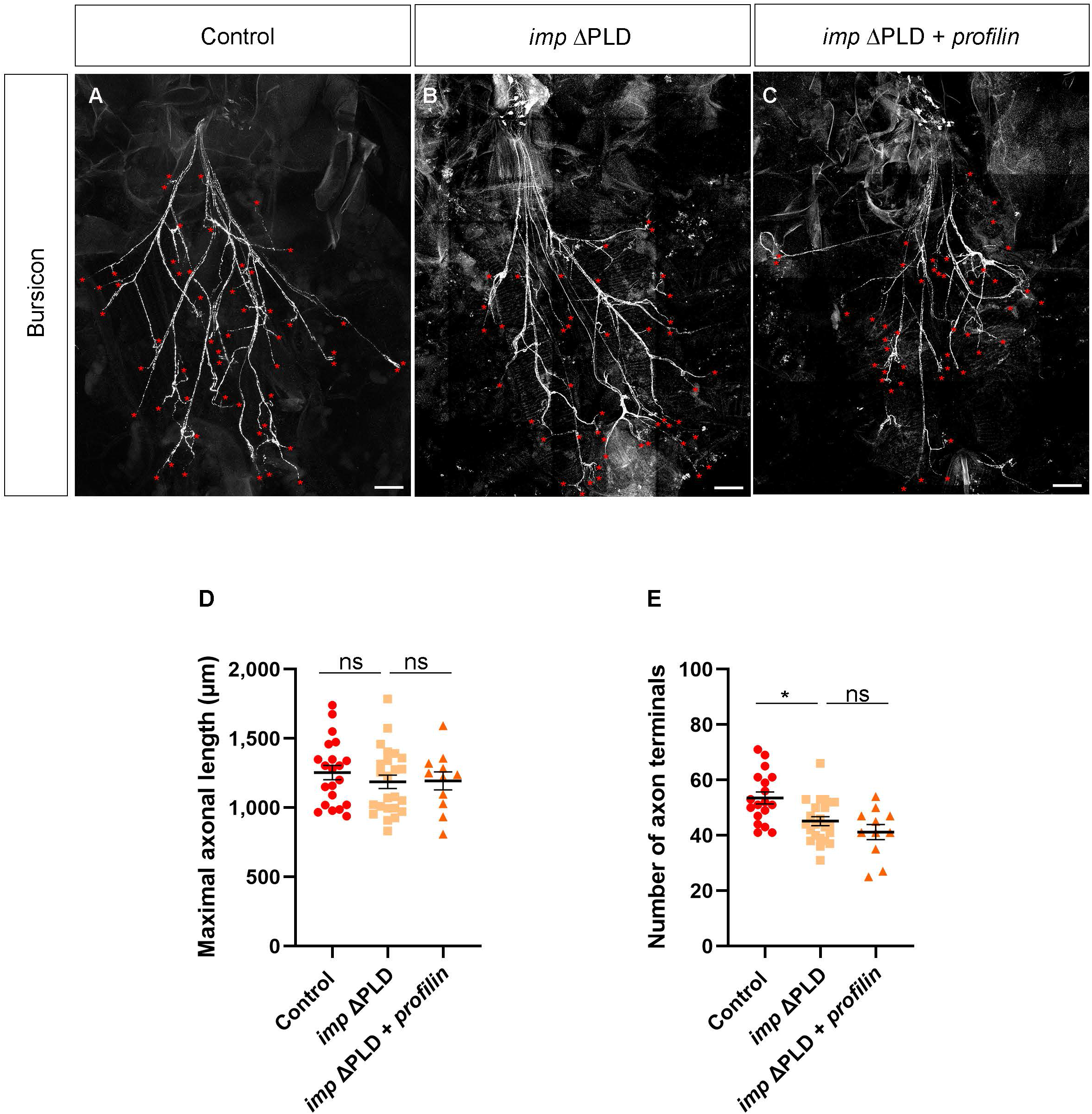
*profilin* overexpression does not rescue CCAP/Bursicon axonal branching defects in *imp* ΔPLD flies. **A-C** Tile confocal images of control (**A**), *imp* ΔPLD (**B**) and *imp* ΔPLD+*profilin* (**C**) whole-mount fillets dissected at 24 h post-eclosion and stained using anti-Bursicon antibody. Red asterisks indicate the axon terminals. Scale bar: 100 µm. Animals were raised at 29°C. Complete genotypes: control: GFP-Imp/Y; CCAP-Gal4,UAS-mCD8-RFP/+; *imp* ΔPLD: GFP-Imp ΔPLD/Y; CCAP-Gal4,UAS-mCD8-RFP/Sco; *imp* ΔPLD+*profilin*: GFP-ImpΔPLD/Y; CCAP-Gal4,UAS-mCD8-RFP/Sco; UAS *profilin*/+. **D** Dot plot showing the maximal axonal length of control, *imp* ΔPLD and *imp* ΔPLD+*profilin* CCAP/Bursicon neurons at 24 h post-eclosion. Number of fillets analyzed: control=21; *imp* ΔPLD=24; *imp*ΔPLD+*profilin*=11. **E** Dot plot showing the number of CCAP/Bursicon axon terminals in control, *imp* ΔPLD and *imp* ΔPLD+*profilin* adult flies at 24 h post-eclosion. Number of fillets analyzed: control=18; *imp* ΔPLD=23; *imp* ΔPLD+*profilin*=11. **D-E** 3 replicates were performed. Quantifications were performed on the same individuals when possible. Bars and error bars correspond respectively to mean and SEM. **P<0.01; *P<0.05 (Kruskall– Wallis with Dunn’s post-tests).

**Supplementary Figure 5:**
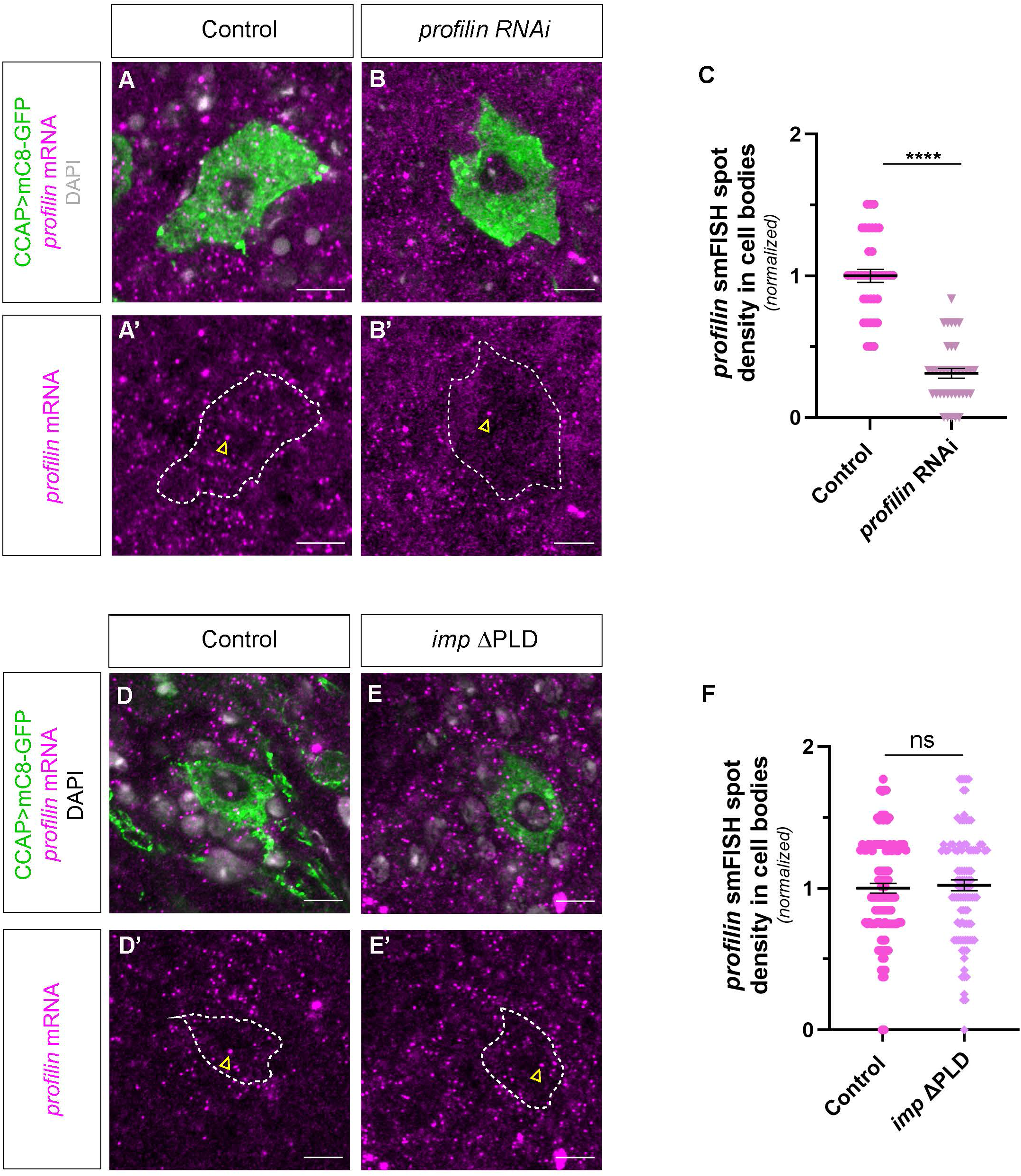
*profilin* mRNA smFISH spot density is not decreased in *imp* ΔPLD CCAP/Bursicon cell bodies. **A**-**B** Confocal images of CCAP/Bursicon neuron cell bodies from control (**A**) and *profilin* RNAi (**B**) whole-mount fillets dissected at 24h post-eclosion and incubated with anti-*profilin* probes. CCAP/Bursicon neurons were detected using CCAP-Gal4 driven mCD8-GFP expression (in green). *profilin* smFISH signals are shown in magenta in **A-B** and **A’-B’**. Brightness and contrast were adjusted for each image for display purposes. Yellow arrowheads point to transcription foci. Scale bar: 5 µm. Animals were raised at 29°C. Complete genotypes: control: Imp-Scarlet/Y; CCAP-Gal4,UAS-mCD8-GFP/+; *profilin* RNAi: Imp-Scarlet/Y; CCAP-Gal4,UAS-mCD8-GFP/UAS-*profilin*-RNAi. **C** Dot plot showing *profilin* smFISH spot density in CCAP/Bursicon cell bodies of control and *profilin* RNAi adult flies at 24 h post-eclosion. Number of CCAP/Bursicon cell body portions analyzed: control=38; *profilin* RNAi=38. 1 replicate was performed. Individual data points correspond to individual regions analyzed (up to 2 per cell body) and were normalized to the control condition. 1 to 5 CCAP/Bursicon cell bodies were analyzed per individual. Bars and error bars correspond respectively to mean and SEM. ****P<0.0001 (two-tailed Mann-Whitney U test on individual data points). **D-E** Confocal images of CCAP/Bursicon neuron cell bodies from control (**D**) and *imp* ΔPLD (**E**) whole-mount fillets dissected at 24h post-eclosion and incubated with anti-*profilin* probes. CCAP neurons were detected using CCAP-Gal4 driven mCD8-GFP expression (in green). *profilin* smFISH signals are shown in magenta in **D-E** and **D’-E’**. Brightness and contrast were adjusted for each image for display purposes. Yellow arrowheads point to transcription foci. Scale bar: 5 µm. Animals were raised at 29°C. Complete genotypes: control: GFP-Imp/Y; CCAP-Gal4,UAS-mCD8-GFP/+; *imp* ΔPLD: GFP-Imp ΔPLD /Y; CCAP-Gal4,UAS-mCD8-GFP/+. **F** Dot plot showing *profilin* smFISH spot density in CCAP/Bursicon cell bodies of control and *imp* ΔPLD adult flies at 24 h post-eclosion. Number of CCAP/Bursicon cell body portions analyzed: control=113; *imp* ΔPLD=101. 3 replicates were performed. Individual data points correspond to individual regions analyzed (up to 2 per cell body) and were normalized to the control condition. 1 to 5 CCAP/Bursicon cell bodies were analyzed per individual. Bars and error bars correspond respectively to mean and SEM. ns stands for not significant (two-tailed Mann-Whitney U test on individual data points).

